# A Large-Scale Structural and Functional Connectome of Social Mentalizing

**DOI:** 10.1101/2020.11.27.400929

**Authors:** Yin Wang, Athanasia Metoki, Yunman Xia, Yinyin Zang, Yong He, Ingrid R Olson

**Author notes:** YW and AM contributed equally to this work. Correspondence: Dr. Yin Wang, Dr. Ingrid Olson.

## Abstract

Humans have a remarkable ability to infer the mind of others. This mentalizing skill relies on a distributed network of brain regions but how these regions connect and interact is not well understood. Here we leveraged large-scale multimodal neuroimaging data to elucidate the connectome-level organization and brain-wide mechanisms of mentalizing processing. Key features of the mentalizing connectome have been delineated in exquisite detail and its relationship with the default mode network has been extensively scrutinized. Our study demonstrates that mentalizing processing unfolds across functionally heterogeneous regions with highly structured fiber tracts and unique hierarchical functional architecture, which make it distinguishable from the default mode network and other social brain networks.

## Introduction

Understanding other people’s intentions, feelings, beliefs, and traits is a pivotal component of human social cognition [1]. This ability is referred to as mentalizing, mindreading, or theory of mind (ToM) [2]. Mentalizing is crucial for successful navigation of the social world, as it allows us to predict, explain, and manipulate each other’s behavior. Atypical mentalizing undermines one’s interpersonal communication, social competence, and overall quality of life [3].

Given its significance for social living, considerable efforts have been made to understand the neural basis of mentalizing [1,4,5]. Existing literature suggests that mentalizing is supported by a widely distributed network of brain regions termed the ‘mentalizing network’ (MTN). The MTN includes the temporoparietal junction (TPJ), precuneus (PreC), anterior temporal lobes (ATL), dorsomedial prefrontal cortex (DMPFC), and ventromedial prefrontal cortex (VMPFC).

The TPJ is believed to be responsible for attributing other’s transient mental states (e.g. instant goals, thoughts, and feelings) [6,7] whereas the DMPFC is thought to be involved in reasoning about other’s enduring traits (e.g. stable attitudes, preferences, and dispositions) [6,8–10]. The PreC has been associated with mental imagery that is necessary for perspective taking (e.g. online mental simulation of how a person thinks, acts or behaves in fictitious situations) [11–14] and portions of the ATL have been shown to be a repository for semantic knowledge related to persons, social concepts, and social scripts [15,16]. Lastly, the VMPFC has been suggested to represent emotional and motivational components of social reasoning (e.g. linking valence to particular persons, their actions, and their thoughts) and exert top-down affective prediction [17–19]. While the precise function of each region is still debated [10,11], they are consistently and reliably recruited for mentalizing, regardless of the task and stimulus formats [8,11,17,20].

Although the functional specialization of single mentalizing area (e.g. TPJ) has been studied intensively, little is known about how these areas are structurally connected and functionally interact. One cause of this gap in knowledge can be ascribed to the revolutionary discovery of the default mode network (DMN) [21]. This prominent brain network has become the primary focus in network neuroscience for years and numerous studies have been carried out to elucidate its anatomy, physiology, functions, and network properties [22]. Since the MTN shows considerable spatial overlaps with the core system of DMN [23–28], some researchers have attempted to assign the DMN’s well-known network properties to the MTN. However, this assumption of ‘connectome resemblance’ has not been systematically investigated and validated. In addition, besides mentalizing, the DMN also shares same set of brain regions with other cognitive functions such as the autobiographical memory [25], moral reasoning [23], self-referential processing [12,29], mental time travel (i.e. remembering the past and imagining the future) [13,30], and semantic memory [31–33]. It is unknown how mentalizing and these seemingly unrelated functions co-exist in the vicinity (neighborhood) of the DMN and how differently each network is structurally organized and functionally operated.

In the present study, we aimed to elucidate and elaborate the full profile of brain connectivity in MTN, such as its network topology, fiber composition, spatial specificity, individual variance, hemispheric preference, structure-function relations, and brain-behavior associations. More importantly, we scrutinized the connectome-level resemblance between the MTN, DMN, and other DMN-vicinity networks. We used the human connectome project (HCP) dataset because it provides a large sample size (n=672) and high quality multimodal neuroimaging data [34]. Due to the heterogeneous nature of cortical areas the MTN encompasses, we functionally defined five bilateral mentalizing regions in each participant (i.e. subject-specific MTN ROIs), then used tractography to delineate local and long-range white matter connections. We also investigated functional and dynamical properties of this network using resting-state and task-state functional datasets. Finally, we identified the nodes of DMN and other functional networks using both meta-analytic and data-driven approaches, and performed analogous tractography and connectivity analyses to compare their connectome characteristics with MTN’s.

## Results

### Structural Connectome

Using probabilistic diffusion tractography techniques [35], we first delineated inter-regional connections between mentalizing areas. Structural connection strength is often interpreted as a measure of capacity for information flow and provides valuable insights about underlying pathways through which neural signals propagate within the network [36]. Here, we reconstructed ten pairwise connections between five mentalizing areas in each hemisphere and characterized the underlying architecture using community detection and hierarchical clustering analysis. We found the organization of structural connectivity (SC) can be divided into two partitions: a lateral subsystem (TPJ-ATL) and a medial subsystem (PreC-DMPFC-VMPFC) (Fig 1).

**Figure 1.**
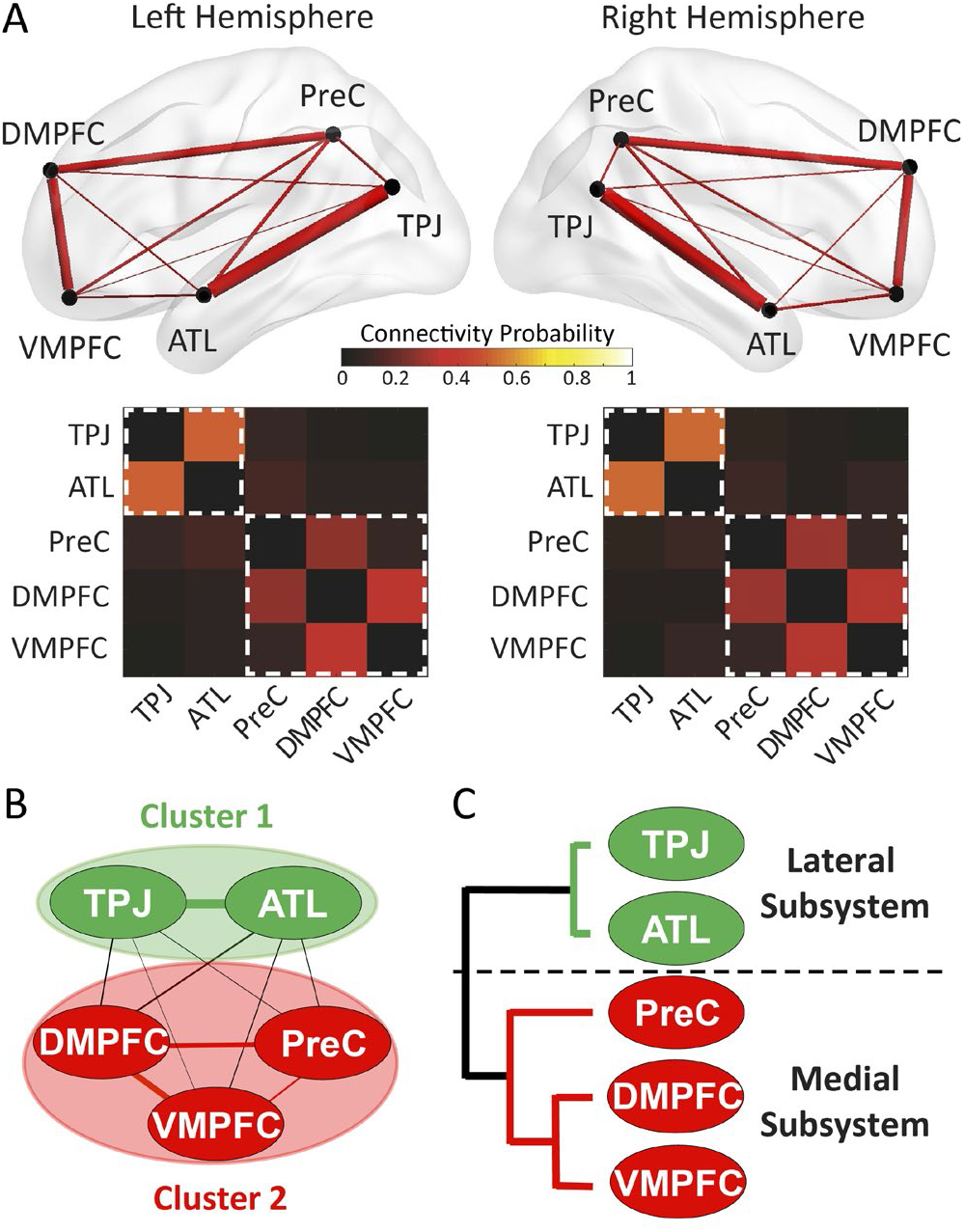
Structural Connectome of the MTN. (A) The pairwise connection strength (that is, connectivity probability) between five MTN ROIs was plotted in wiring diagrams (top) and weighted matrices (middle). The width of each line in the wiring diagrams (or the heat color in the matrices) proportionally reflects the mean connectivity probability across all individuals. The landscape of structural connectivity maps clearly indicated two partitions: a medial subsystem (PreC-DMPFC-VMPFC) and a lateral subsystem (TPJ-ATL). Further analyses using community detection algorithm (B) and hierarchical clustering algorithm (C) confirmed this medial-lateral division. Highly similar connectivity patterns and partitions were also obtained when using another SC measure (fiber count maps) (i.e. correlations between these two maps in LH: r(10) =0.995, p<0.001, 95% CI: 0.916, 1.074; and in RH: r(10) =0.996, p<0.001, 95% CI: 0.923, 1.069).

Next, we investigated the white matter composition of the MTN by dissecting its constituent fiber tracts. A small number of long-range tracts have been repeatedly reported in the literature and disruption of these tracts can lead to ToM impairments [37]. Lesion samples are naturally heterogeneous, including individuals with small and large lesions, as well as lesions that destroy differing amounts of gray matter and white matter. Thus, it is important to extend this line of research to a neurologically normal sample and investigate at a higher granularity (i.e. quantify the fiber composition of each MTN connection). Here we used an automatic fiber reconstruction technique [38] to delineate ten major fasciculi for each subject and compared them with the ROI-ROI tracts at the individual level. By calculating the degree of their overlapped trajectories, we can estimate the involvement of major fasciculi in the mentalizing connectome. In agreement with prior meta-analysis [37], our results revealed five major fasciculi (Fig 2A). Specifically, precuneus-related connections were primarily constructed by the cingulum bundle (CING, e.g. PreC-DMPFC, 37±19%, (mean ± SD)); ATL-related connections were mainly supported by the inferior longitudinal fasciculus (ILF, e.g. ATL-TPJ:37±16%, ATL-PreC:31±12%); TPJ-related connections were largely overlapped with the superior longitudinal fasciculus and arcuate fasciculus (SLF/AF, e.g. TPJ-DMPFC: 20±10%), and VMPFC-related connections were mainly mediated by the inferior fronto-occipital fasciculus (IFOF, e.g. VMPFC-TPJ: 29±13%) and uncinate fasciculus (UF, e.g. VMPFC-ATL: 28±16%). As a whole, the medial subsystem appears to be solely subserved by the cingulum bundle, whereas the lateral subsystem is supported by multiple white matter bundles (ILF and SLF/AF), and the two subsystems are linked by the UF, IFOF, and SLF/AF.

**Figure 2.**
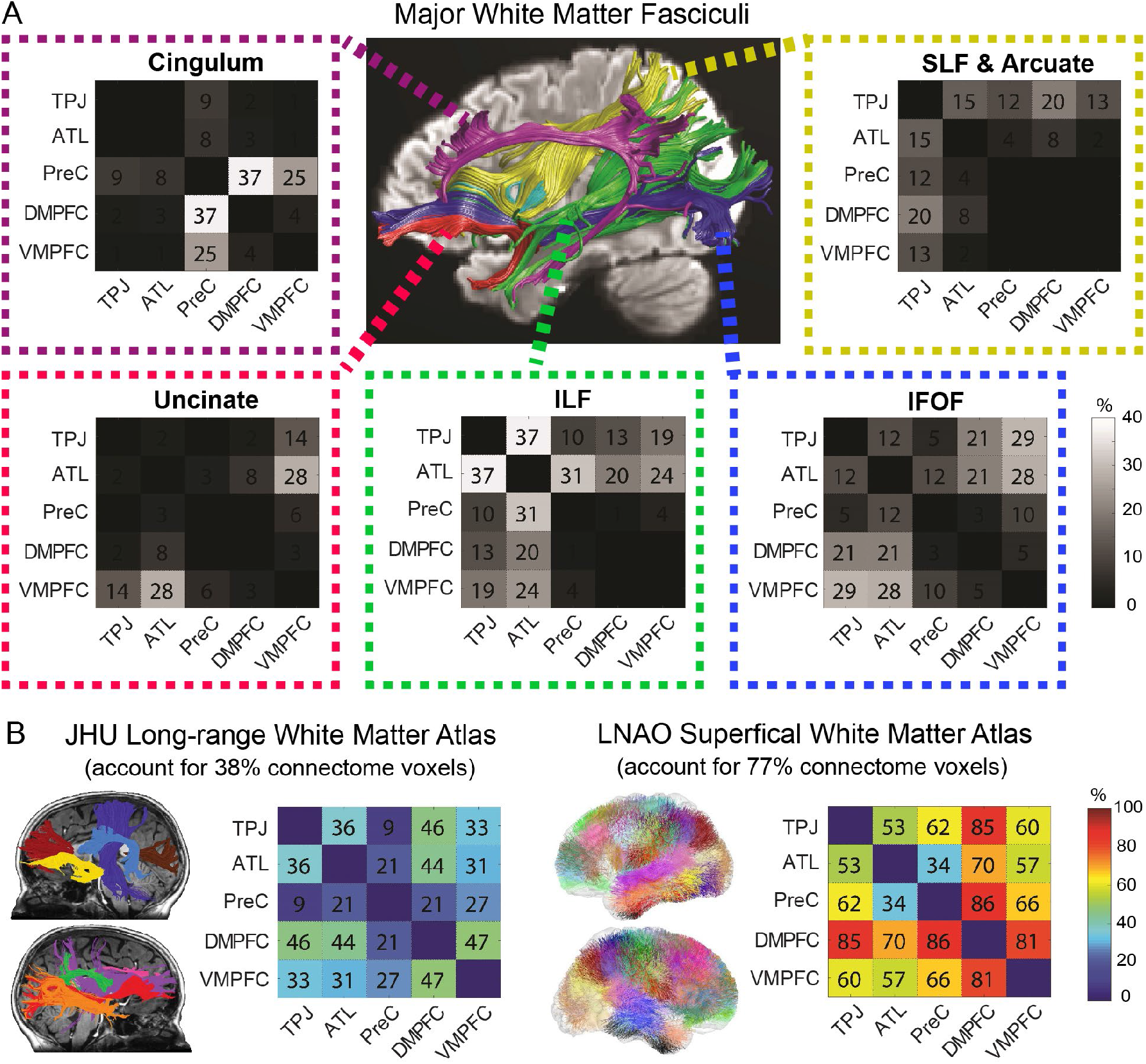
White Matter Composition of the MTN. (A) Top five major fasciculi that overlap mostly with the mentalizing connectome. Each percentage number in the matrices represents the mean overlapped volumes between an ROI-ROI connection and a major fasciculus across all subjects. Taking the TPJ-VMPFC connection as an example, 29% voxels of the tract overlapped with the IFOF and 19% overlapped with the ILF; the tract did not overlap with the cingulum (0%) and had small portion with the SLF (13%) and uncinate (14%). (B) The relative contribution of large-range and short-range fibers to the mentalizing connectome. After overlaying all ROI-ROI connections onto two standard white matter atlases, we found ~77% of the MTN connectome can be classified as short-range fibers whereas only ~38% can be labelled as long-range fibers. For each ROI-ROI connection, the numbers in the matrices represent how well it can be explained by the tracts in each atlas. Taking the TPJ-VMPFC connection as an example, 33% of its voxels overlapped with long-range tracts in the JHU atlas whereas 60% of voxels overlapped with superficial tracts in the LNAO atlas. Note that human white matter is somewhat idiosyncratic across individuals and the two standard atlases only represent the skeleton of the most common white matter fibers in standard MNI space. Since they were created by different research groups with different methods, the two atlases are not mutually exclusive and their combination does not always explain all connectome voxels (e.g. the sum of PreC-ATL by two atlases is only 55%). All numbers here were averaged across two hemispheres (see Supplementary Fig 1 for details).

Since only a small proportion of the connectome can be attributed to long-range fasciculi in deep white matter (e.g. the highest percentage in Fig 2A was 37%), we directed our attention to another important class of white matter—short-range superficial fibers (also called U-shaped fibers). Tracer studies have demonstrated that long-range fibers comprise only a minority of the whole brain connectome with the majority consisting of short association fibers that lie immediately beneath the gray matter, connecting adjacent gyri [39,40]. By projecting all ROI-ROI connections over two standard white matter atlases of long-range [41] and short-range fibers [42], we interrogated the relative contribution of the two white matter systems for the mentalizing connectome. While only 37.64% of the connectome can be labelled by the fasciculi in the long-range white matter atlas, more than 77.02% of the ROI-ROI connections were classified as short-range fibers (Fig 2B). Moreover, we found each ROI-ROI connection was supported redundantly by both types of fibers but that the short-range ones were more involved than the long-range ones.

### Functional Connectome

To understand how mentalizing areas functionally interact with each other, we examined their inter-regional functional connectivity (FC) across different brain states and tasks. The HCP dataset provides multiple in-scanner cognitive tasks requiring different degrees of mentalizing. For example, the HCP ToM task explicitly asks subjects to implement mental state attribution when watching Heider and Simmel-type social animations. The HCP emotion task requires subjects to recognize and understand the emotions displayed in facial expressions and thus elicits implicit mentalizing [9,43]. Lastly, the HCP motor task asks subjects to execute simple body movements (fingers, toes, or tongues) and thereby engages minimal mentalizing. As showed in Fig 3A, we found highly similar FC topological patterns across three HCP task categories (all Pearson’s *rs*>0.342; t-tests: *ts*>26.85, *p*s<0.001, see details of pairwise pattern similarities between task categories in Supplementary Table 2a), suggesting the existence of an ‘intrinsic’ coherent functional architecture that is constantly active and synchronized across contexts [44–46]. This intrinsic network architecture was also confirmed by using the HCP resting state data (Fig 3B). Moreover, the strength of FC seems to depend on the cognitive state, with elevated FC when the task requires more mentalizing (i.e. motor-state FC < emotion-sate FC < ToM-state FC). Statistical testing using multilevel modeling analysis further confirmed this effect by showing that all ten pairwise FCs in MTN followed the same trend, that is, the degree of mentalizing required by a task significantly predicted the strength of FC elicited by that task category (Supplementary Table 2b). To further validate its specificity to the MTN, we applied analogous cross-tasks FC analyses to another brain network (i.e. the DMN) but found no such progressive effect (Supplementary Fig 6). It should be noted that we were not able to directly compare the size of FCs between the resting-state and three task-states as they had substantial differences in scan length (i.e. 56mins for resting versus 6 mins for cognitive tasks).

**Figure 3.**
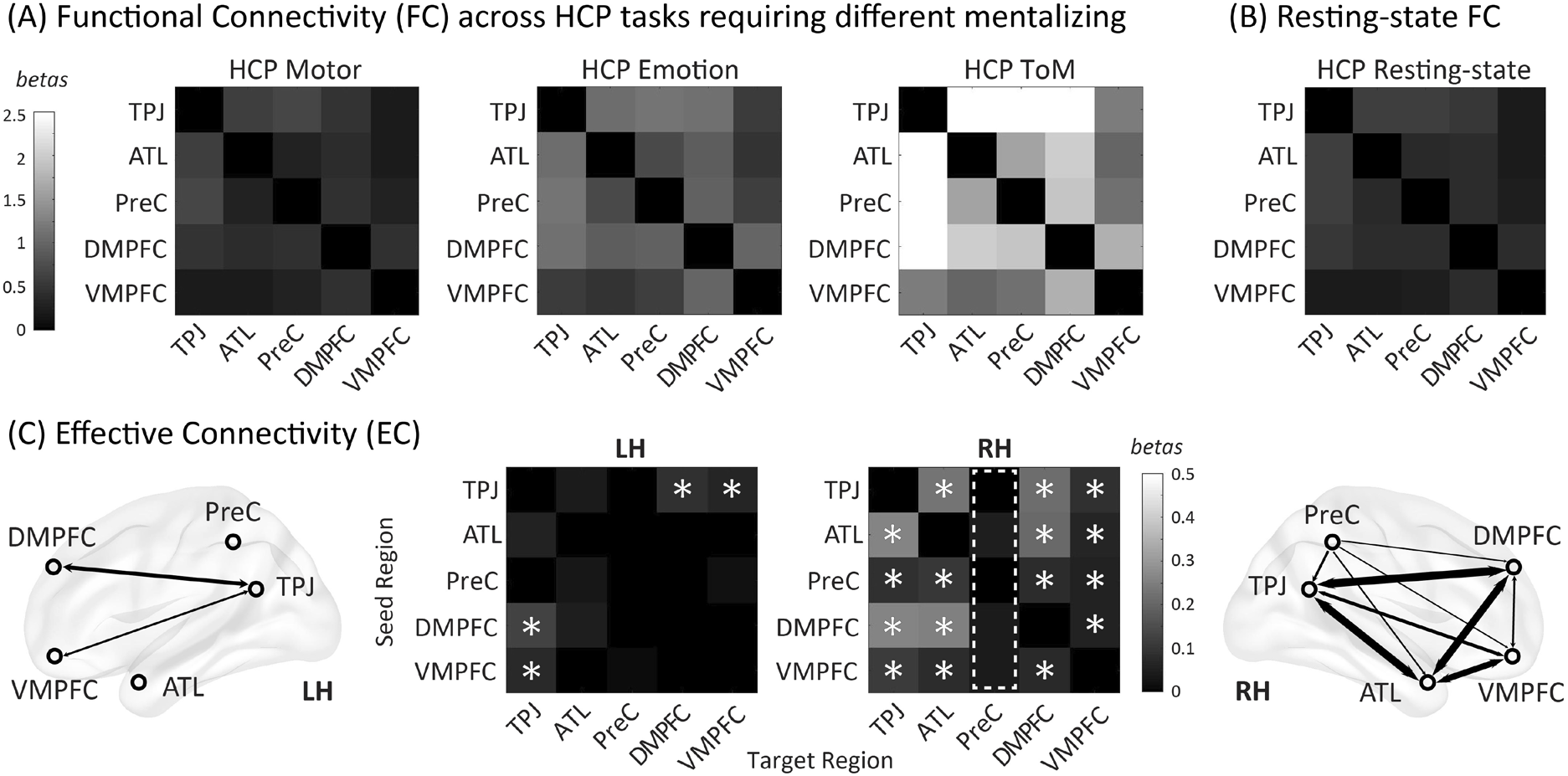
Functional Connectome of the Mentalizing Network. (A) Functional connectivity (FC) in MTN across different cognitive tasks. We selected three HCP tasks with different degrees of mentalizing: the motor (no mentalizing), emotion (implicit mentalizing), and ToM task (intentional mentalizing). We found similar FC topological patterns across tasks, with high FC consistently in TPJ-related and DMPFC-related connections. Intriguingly, coactivations among mentalizing areas were dependent on the degree of mentalizing required in the task: when there were increased mentalizing demands, FC increased gradually from the motor-state, emotion-state, to ToM-state task. We only show the right hemisphere matrices here, but the left side had the same effect. (B) FCs among MTN areas in the resting state. Note that the topological patterns in resting state were highly similar with three HCP tasks, indicating the existence of an ‘intrinsic’ coherent functional architecture that is constantly synchronized across brain states. (C) Effective connectivity during the HCP ToM task. PPI matrices for each hemisphere were showed in the middle (seed regions in y-axis and target regions in x-axis; asterisks indicate statistically significance) and an illustration of directional information flow summarized from the matrices were added on two sides (arrow line width was scaled to reflect the EC strength). The two hemispheres exhibited entirely different EC patterns, with much stronger task-modulated dynamics in the right hemisphere. For the right hemisphere, the precuneus was identified with only feed-forward information cascades (the dashed box indicates the non-significant feed-back processing toward precuneus), whereas other four mentalizing areas implicate recurrent processing during mentalizing. We did not find any significant EC among MTN ROIs in the HCP motor or emotion task.

Next, we investigated the organization of the intrinsic functional architecture, as it can provide insights about how information is functionally integrated within the MTN. Only one unified community was detected (i.e. no modularity), which is different from the two parallel subsystems we found in the SC analysis. Hierarchical clustering analysis, however, identified a consistent functional hierarchy across all resting and task FC maps—with TPJ and PreC as the lowest level and VMPFC as the highest level (see Supplementary Fig 3B). This organization implies a serial-hierarchical functional architecture in which information integrates from posterior MTN ROIs to anterior ROIs and corresponds well with the literature on the functional role of each mentalizing area [11].

As FC only expresses statistical dependencies (i.e. non-directional correlations) among time courses of different brain areas [36], we also examined the effective connectivity (EC) which allows us to capture stimulus-driven patterns of directional influence among mentalizing areas [47]. Psychophysiological interaction (PPI) analyses build simple static models of EC between one or more brain regions and allow researchers to explore directed changes in connectivity by establishing a significant interaction between the seed region and the psychological context [47–49]. Our PPI analyses revealed imbalanced information processing across the hemispheres, with much stronger task-modulated brain dynamics in the right hemisphere than the left hemisphere (Fig 3C). In addition, the precuneus in the right hemisphere was found to be only engaged in feed-forward processing, whereas other mentalizing areas had both feed-forward and feed-back EC during social mentalizing.

### Connectome Features: Individual Variance

Previous studies showed striking individual differences in ToM task performance [50–52]. Here we examined whether such variance can be observed in the connectome. To reveal the distribution of group-to-individual similarity across all subjects, we conducted correlation analyses between group-averaged maps and subject-specific maps for each type of brain connectivity measure (see histograms and stats in Supplementary Fig 2A). Results suggested that group-averaged matrices were strongly correlated with subject-specific matrices for SC, rsFC and ToM-state FC (all mean *rs* > 0.424; *ts* > 35.33, *ps* < 0.001, n=672) and majority of subjects (>52.3%) exhibited highly similar maps (r > 0.5) as the group-averaged ones. The group-averaged EC matrices, however, showed mild correlation with subject-specific EC matrices (all mean *rs* > 0.136; *ts* > 9.67, *ps* < 0.001, n=672) and most subjects (>57.2%) exhibited small correlation (r > 0.1) with the group-averaged ones.

Next, we used two cross-validation schemes to further probe subgroup-to-subgroup and group-to-individual similarity, that is, the split-half cross-validation (using 50% subjects’ matrices to predict the other 50% subjects’ matrices) and the leave-one-out cross-validation (using N-1 subjects’ matrices to predict a new subject’s matrices) (see Supplementary Fig 2B). Again, both methods revealed high train-test correlation for SC, rsFC and ToM-state FC (all mean *rs* > 0.419; *ts* > 35.08, *ps* < 0.001, n=1000/672) and relatively small train-test correlation for the EC (all mean *rs* > 0.124; *ts* >9.55, *ps* < 0.001, n=1000/672). Taken together, the brain connectivity in MTN was found to be homogeneous across subjects, with large homogeneity in SC, medium in FC, and small in EC. Thus, our reported group-averaged matrices indeed reflect most individuals’ connectivity patterns. We also compared the individual variance of MTN with other brain networks and found weaker group-to-individual similarity in MTN than the DMN (Supplementary Table 6) and the face processing network [53] for all brain connectivity types. This suggests that MTN is relatively a more heterogeneous network.

### Connectome Features: Spatial Specificity

To explore how connectivity patterns are spread and changed in space, we performed analogous tractography and connectivity analyses while manipulating the methods of ROI location selection. Past research has documented three different ways to define the spatial location of ROIs, either using subject-specific coordinates from a functional localizer, or group-level peak activation from a task, or putative coordinates from meta-analyses [54,55]. Here we examined how these precise or liberal methods impact MTN connectivity maps. In addition, since remarkable individual variability has already been observed in our subject-specific coordinates (see black dots in brain images in Fig 4), we also examined how shuffling subject-specific ROIs across subjects (so that the TPJ coordinates of subject A became the TPJ coordinates of subject B) alters connectivity maps. The results revealed that the functional connectome (rsFC, ToM-state FC and EC) was very susceptible to ROI selection whereas the structural connectome remained stable across all location manipulations (Fig 4). Although connectivity maps using liberal methods were still statistically correlated with the original subject-specific maps (SC: all mean *rs*>0.62, *ts*>72.75, *ps*<0.001; rsFC: all mean *rs*>0.24, *ts*>5.50, *ps*<0.001; ToM-state FC: all mean *rs*>0.21, *ts*>5.34, *ps*<0.001; EC: all mean *rs*>0.08, *ts*>2.49, *ps*<0.013), fine-grained connectivity patterns was only preserved in SC but systematically changed in rsFC, ToM-state FC and EC. Of most interest, adopting more liberal methods generated connectivity maps that were more dissimilar to the original subject-specific ones (Fig 4), as pattern similarity decreased substantially from methods using group-level peak activation, to methods using Neurosynth coordinates, and to methods using shuffled coordinates (i.e. the drop of Pearson’s r in SC (0.70→0.65→0.62), rsFC (0.34→0.26→0.24), ToM-state FC (0.34→0.28→0.21), EC (0.11→0.09→0.08)). Overall, these findings provide evidence of high spatial specificity in the functional connectome but low spatial specificity in the structural connectome.

**Figure 4.**
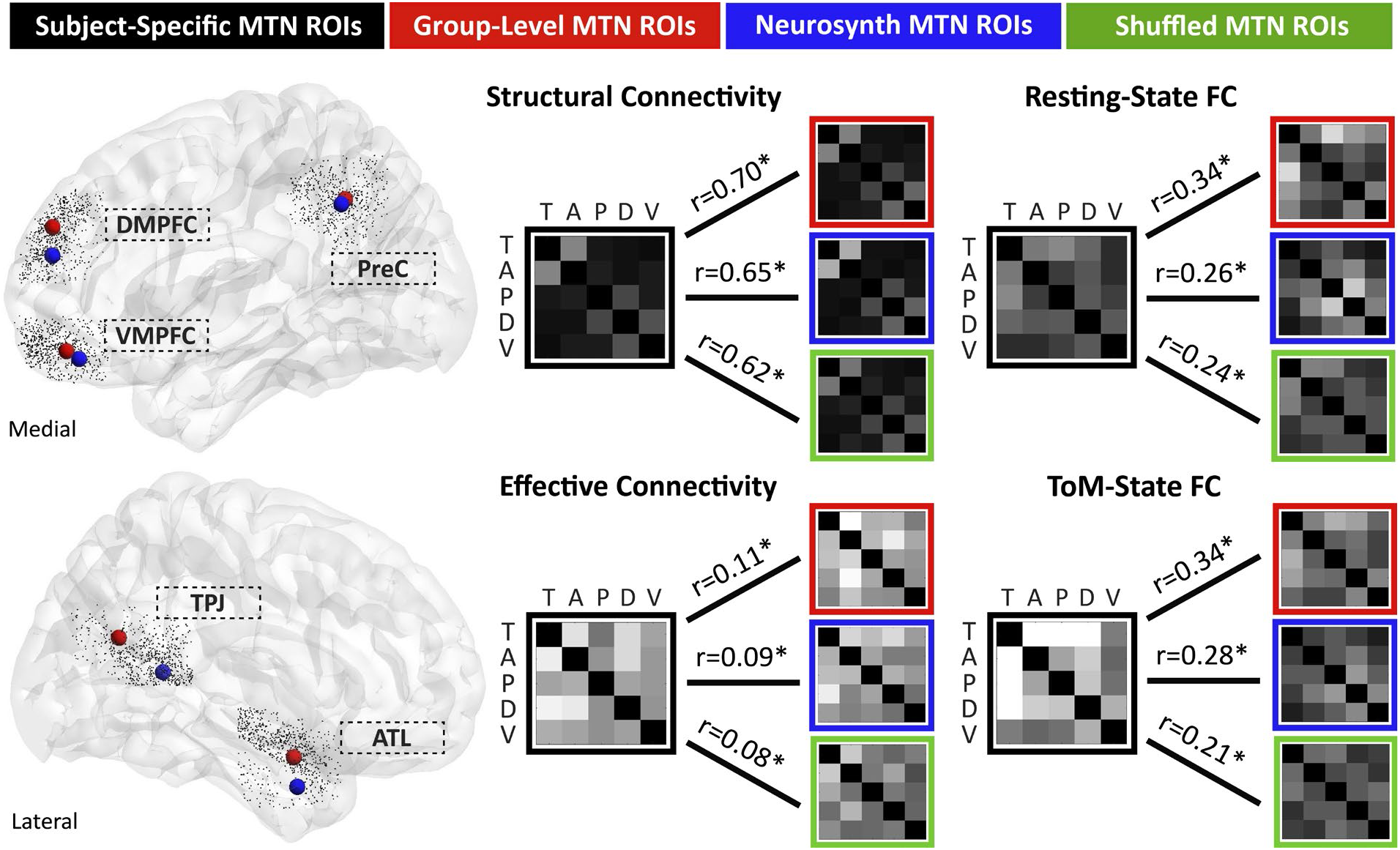
The Spatial Specificity of the Mentalizing Connectome. Brain connectivity patterns were measured and compared across four different ways of ROI selection. We originally defined each mentalizing ROI by using subject-specific coordinates from the functional ToM localizer. The black dots in left-side brain images showed the massive individual variability within each node (spatial distributions in TPJ (14mm), ATL (13mm), PreC (12mm), DMPFC (11mm), and VMPFC (11mm)). The major findings in this article came from this method (black frames). We then adopted two more liberal ways to define ROI locations, by choosing either group-level peak activation sites from the functional ToM localizer (red dots and frames) or putative coordinates from Neurosynth database (blue dots and frames). Lastly, we permuted subject-specific ROI locations across subjects (e.g. subject A’s DMPFC coordinates now became subject B’s) and defined ROIs with shuffled coordinates (green frames). We found the functional connectome (rsFC, ToM-state FC and EC) was very susceptible to spatial location changes because all fine-grained patterns in the original subject-specific maps were substantially altered, whereas the SC maps remained stable after location manipulations and they were all highly correlated with original subject-specific maps (all mean *rs*>0.6). Critically, the more liberally one chose to define ROIs (red→blue →green), the maps were more dissimilar to the original subject-specific maps. This emphasizes the importance of precise ROI definition when studying fine-grained connectivity fingerprints in MTN. For simplicity, we only provided the connectivity maps in the right hemisphere here but the results were very similar in the left hemisphere. Asterisks indicate statistical significance (p<0.05). Abbreviations: T=Temporo-parietal junction (TPJ); A=Anterior Temporal Lobe (ATL); P=Precuneus (PreC); D=Dorsal Medial Prefrontal Cortex (DMPFC); V=Ventral Medial Prefrontal Cortex (VMPFC)

### Connectome Features: Functional Specificity

The MTN shares similar nodes with the core system of DMN as well as other functional networks in the vicinity of DMN (i.e. autobiographical memory, moral reasoning, self-reference, mental time travel, and semantic memory) [12,23,32]. Here we defined these networks by using their Neurosynth coordinates (see Supplementary Table 4) and performed analogous connectivity analyses to reveal their connectome resemblance with the MTN (Fig 5 and Supplementary Fig 4). We found the MTN had highly similar SC patterns with the DMN and nearby networks (all mean *rs*>0.47, *ts*>12.60, *ps*<0.001) and they were all organized by the same structural architecture (‘lateral vs medial’ subnetworks) except the semantic memory network (Fig 4). This is consistent with our results in the spatial specificity analysis showing that the SC was configured as relatively stable across spatial extent. For FC (rsFC and ToM-state FC), despite the significant global correlations between the MTN, DMN and nearby networks (all mean *rs*>0.15, *ts*>5.82, *ps*<0.001), they all exhibited distinct local connectivity patterns. For EC, even the global pattern correspondence did not exist among all networks (all mean *rs*<0.07, *ts*<1.77, *ps*>0.077), let alone their disparate fine-grained connectivity fingerprints. Therefore, our results revealed that all DMN-vicinity networks share a similar anatomical architecture (fiber bundles) but own distinguishable functional and dynamic characteristics (in particular the EC).

**Figure 5.**
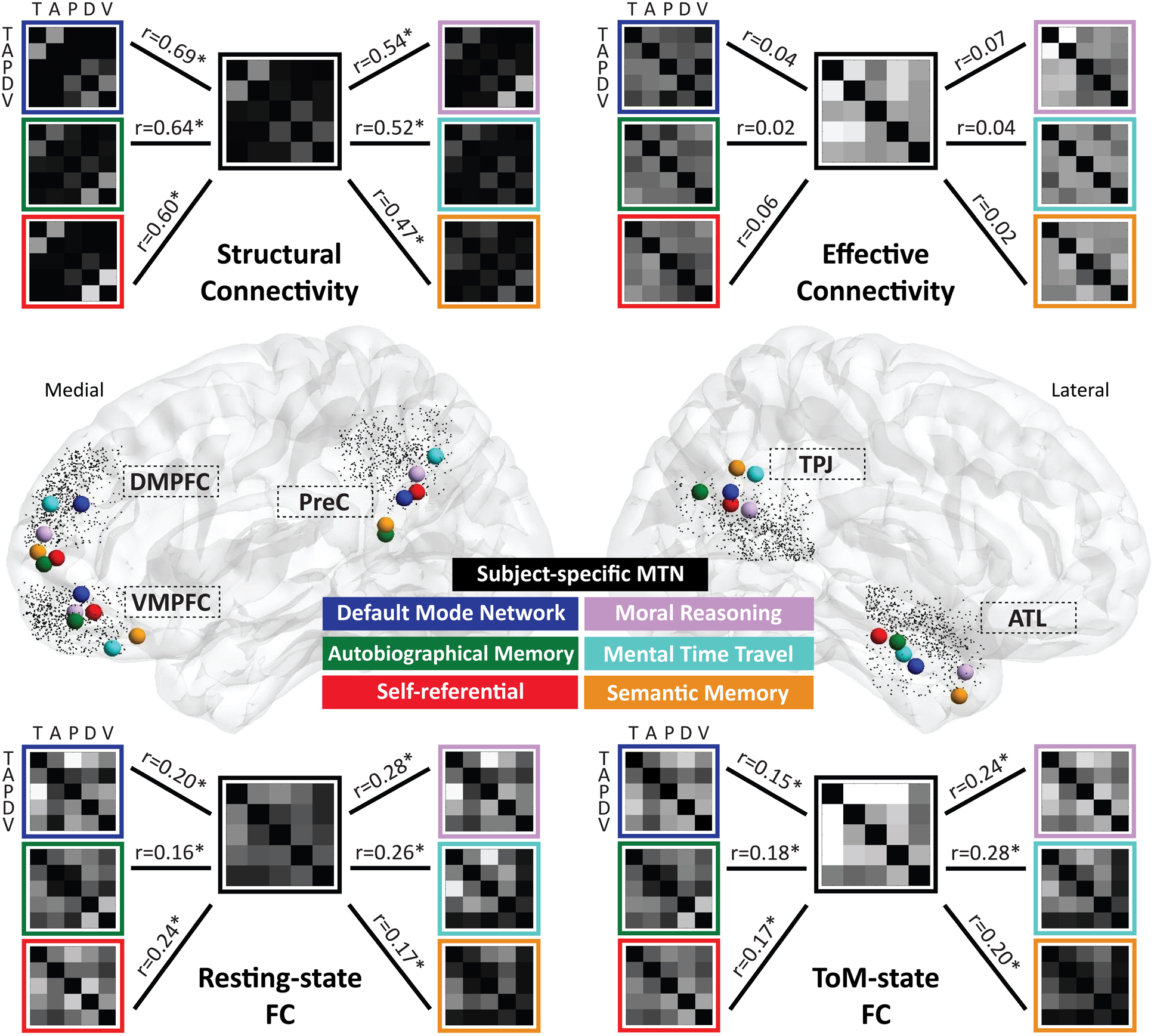
Brain connectivity similarity between mentalizing network, default-mode network, and other nearby functional networks. Brain connectivity patterns were measured and compared across seven DMN-vicinity networks (node locations are depicted in different colors). The MTN was defined using subjects-specific coordinates from the HCP ToM task. Six other networks were defined using coordinates from Neurosynth (DMN, autobiographical memory, self-referential, moral reasoning, semantic memory) or meta-analysis (mental time travel) (see search keyword and specific coordinates in Supplementary Table 4). Tractography analyses (upper left panel) revealed highly similar SC patterns between all networks (all mean *rs*>0.47, *ps*<0.001) and they (except semantic memory) were all organized by the same architecture with two subsystems (i.e. medial vs lateral). For resting-state and ToM-state FC (two lower panels), the global pattern of all networks was statistically correlated (all mean *rs*>0.15, *ps*<0.001) but each showed distinct local fine-grained FC. For EC (upper right panel), all networks exhibited different global and fine-grained patterns (correlations were all insignificant, *rs*<0.007, *ps*>0.77), indicating that each DMN-vicinity network had unique dynamic connectivity properties. Asterisks indicate statistical significance (p<0.05). Abbreviations: T=Temporo-parietal junction (TPJ); A=Anterior Temporal Lobe (ATL); P=Precuneus (PreC); D=Dorsal Medial Prefrontal Cortex (DMPFC); V=Ventral Medial Prefrontal Cortex (VMPFC)

To complement our meta-analytic approach that imperfectly assumed all subjects had the same DMN node locations (i.e. Neurosynth), **we also validated the findings by adopting a data-driven method (i.e. spatial ICA) to identify subject-specific DMN** [56]. **A similar relationship was observed between personalized MTN and personalized DMN** (i.e. similar SC but dissimilar FC and EC; see Supplementary Fig 5).

### Connectome Features: Lateralization, Structure-Functional Relation, and Brain-Behavior Association

Our original plans did not include conducting a laterality analysis. However, since the PPI analyses showed remarkably unbalanced task-modulated connectivity dynamics across two hemispheres, we subsequently examined hemispheric lateralization at all levels of the connectome. Repeated-measure ANOVA revealed a significant main effect of hemispheric asymmetry in task-evoked BOLD response in mentalizing ROIs (F(1,671) = 109.45, p<0.001, η2 = 0.582) and EC (F(1,671) = 39.74, p<0.001, η2 = 0.056). Despite the absence of a main effect of hemispheric asymmetry, we found that a significant interaction between hemispheric asymmetry and pairwise connections existed in SC (F(9,6039) = 11.05, p<0.001, η2 = 0.016), rsFC (F(9,6039) = 4.53, p<0.001, η2 = 0.007) and ToM-state FC (F(9,6039) = 4.91, p<0.001, η2 = 0.007). This interaction suggests that different pairwise connections might have different directions of lateralization. Post-hoc t-tests further revealed that most asymmetrical connections were right lateralized, indicating that the MTN is a right hemisphere dominated network (see Supplementary Table 3a). In contrast, the DMN appears to be left hemisphere dominated. Repeated-measure ANOVA revealed significant main effect of hemispheric asymmetry in DMN’s SC (F(1,671) = 549.40, p<0.001, η2 = 0.450) and rsFC (F(1,671) = 670.38, p<0.001, η2 = 0.500) and most asymmetrical connections were left lateralized (see Supplementary Table 3b)

We also examined the structure-function relationship within the MTN (Supplementary Fig. 3). Across the entire network, SC and FC were globally correlated with each other (right SC & rsFC: mean r=0.200, t(671)=16.78, p<0.001, d=0.65, 95% CI: 0.177, 0.223; right SC & ToM-state FC: mean r=0.170, t(671)=14.56, p<0.001, d=0.56, 95% CI: 0.147, 0.193; left SC & rsFC: mean r=0.170, t(671)=13.80, p<0.001, d=0.53, 95% CI: 0.146, 0.195; SC & left ToM-state FC: mean r=0.168, t(671)=14.39, p<0.001, d=0.56, 95% CI: 0.145, 0.191). This supports the idea that FC is generally dictated by the underlying white matter [36,57]. However, when inspecting each connection, we found SC and FC did not always correspond at a fine-scale. For example, TPJ had strong FC with PreC and DMPFC but exhibited weak SC with both areas (Supplementary Fig 3A). These disagreements suggest that FC is a good but imperfect representation of the underlying SC [58,59]. In addition, EC maps were found to be neither similar to SC (EC & SC: mean r=0.009, t(671)=0.74, p=0.459, d=0.03, 95% CI: −0.014, 0.031) nor FCs (EC & rsFC: mean r=0.024, t(671)=1.81, p=0.071, d=0.07, 95% CI: −0.002, 0.050; EC & ToM-state FC: mean r=0.022, t(671)=1.64, p=0.102, d=0.06, 95% CI: −0.004, 0.049), owing to its nature of reflecting asymmetrical and directional information processing during mentalizing. Finally, hierarchical clustering and graph theory analysis uncovered more organizational differences between SC and FC: while the structural architecture of MTN is configured by two parallel subsystems with one hub (PreC), the functional architecture is organized by a serial-hierarchical structure with two functional hubs (TPJ and PreC) (Supplementary Fig 3B and Table 5). Overall, we found the SC and FC in MTN were tightly linked but each had their own unique properties.

Finally, we explored if any connectome features were associated with individual’s ToM task performance. Pearson correlations revealed that accuracy on the HCP ToM task was significantly correlated with neural responses in bilateral TPJ (right TPJ: r(672)=0.134, p<0.001, 95% CI: 0.059, 0.209; left TPJ: r(672)=0.184, p<0.001, 95% CI: 0.109, 0.258), bilateral ATL (right ATL: r(672)=0.159, p<0.001, 95% CI: 0.084, 0.234; left ATL: r(672)= 0.126, p=0.003, 95% CI: 0.050, 0.201), and bilateral DMPFC (right DMPFC: r(672)=0.097, p=0.020, 95% CI: 0.021, 0.172; left DMPFC: r(672)=0.120, p=0.004, 95% CI: 0.045, 0.195). While the task accuracy cannot be predicted by any features of SC (all *rs* < 0.085, *ps* > 0.280), rsFC (all *rs* < 0.106, *ps* > 0.080) and ToM-state FC (all *rs* < 0.105, *ps* > 0.075), it was significantly associated with three connections in EC: right ATL→TPJ (r(672)=0.136, p<0.001, 95% CI: 0.061, 0.211), right TPJ→ ATL (r(672)=0.108, p=0.044, 95% CI: 0.033, 0.184), and right DMPFC→TPJ (r(672)=0.121, p=0.040, 95% CI: 0.046, 0.196). These findings highlight the usefulness of dynamic properties of the connectome (i.e. local task-evoked responses in right ATL and TPJ as well as their task-modulated connectivity dynamics) in explaining individual variance of mentalizing skills, whereas those stable connectome features reflecting intrinsic architecture (i.e. SC and FC) cannot. No brain features were found to correlate with the ToM task speed (all |rs|<0.086; all ps>0.078).

## Discussion

The neural basis of theory of mind has been investigated for more than 25 years [60]. While much work has focused on the functional segregation in the MTN (i.e., specific function operated by single mentalizing area) [11], the connectome-level organization and brain-wide mechanisms for functional integration of mentalizing processing remains poorly understood. New trends in connectomics suggest that the function of any mentalizing areas should be considered using an integrative approach, including not only patterns revealed by local properties, but also connectivity and dynamics with other mentalizing areas [61]. Prior connectivity studies on MTN, however, were methodologically limited by small sample size [62–64], the examination of only partial connectomes [65], non-multimodal approaches that neglected white matter connectivity [37], or a restricted focus on atypical groups such as individuals with clinical disorders [37,66,67]. On the other hand, network neuroscience traditionally uses large multimodal data to characterize the entire human connectome or large-scale domain-general networks (e.g. DMN, salience network, frontoparietal control network) but pays less attention to functionally specialized networks with small numbers of nodes and edges. The present study bridges these two domains by elucidating the full profile of brain connectivity in the MTN in exquisite detail and by systematically assessing the relationship between domain-general and domain-specific brain networks (DMN vs. MTN). **For overall readability and accessibility, all methods, analyses and findings in the present study are summarized in** **Supplementary Table 7**.

First, we investigated the anatomical architecture of MTN. We found that the “wiring system” –fiber tracts - was characterized by two separate partitions: a lateral subsystem consisting of the TPJ-ATL and a medial subsystem consisting of the PreC-DMPFC-VMPFC. There are multiple ways to understand this medial-lateral configuration. It has been well-documented that cortical midline areas support self-representation and self-processing [12,68–70] whereas lateral areas are involved in retrieving knowledge about other entities (e.g. ATL) [15,71] and reasoning about others’ perspectives (e.g. TPJ) [11,72]. Given that meta-representation (i.e. simultaneous representation of one’s own and other’s mental states) and decoupling (i.e. self-other distinction) are such essential ingredients for mentalizing, the medial and lateral subsystem may subserve separate pathways for self (egocentric) and other (altercentric) processing. Another interpretation is that these two subsystems might be recruited for different types of ToM inference. It has been reported that lateral regions (e.g. TPJ) are recruited more by stimuli and tasks requiring reasoning about others’ beliefs and intentions (epistemic or “cognitive” ToM) while medial regions (e.g. VMPFC) are recruited more by inferences about emotions and preferences (motivational or “affective” ToM) [18]. Note that it is important to consider whether this partition is a misleading by-product from the limited tractography technique, since fiber tracking algorithms don’t work well when they have to travel through deep white matter from a lateral to medial direction. Future research should test these ideas to clarify the function of each MTN subsystem.

Second, we examined the white matter composition of the MTN at a high granularity level. We found that local U-shaped fibers play a disproportionate role in the scaffold of the MTN. Based on a recent literature review [37], five long-range fiber bundles—the cingulum, SLF, UF, IFOF and ILF—are the most frequently reported tracts associated with theory of mind; in contrast, local white matter was rarely mentioned. In addition, since most MTN ROIs are spatially far apart (e.g. 80% connections had Euclidian distance > 70mm), long-range fibers would be expected to play the primary role in interregional connections. However, our results suggest that long-range fiber bundles only account for 38% of the connectome, whereas short-range fibers support most ROI-ROI connections. This observation is consistent with older histology studies showing that the human cerebral white matter is dominated by short-range fibers that connect adjacent gyri [39] and complies well with the ‘small-world’ characteristic of brain networks [73]. It is believed that abundant short-range fibers serve to adaptively minimize global wiring costs, whereas sparse long-range connections contribute to functional integration [74,75]. Our analysis revealed that most ROI-ROI connections were redundantly constructed by both types of fibers (Fig. 2B). For example, the TPJ and DMPFC are directly linked by major fasciculi, but they are also connected indirectly by local U-shaped fibers via multiple-hop-relays. A redundant wiring scheme can provide resiliency in the case of brain injury or disease, which may explain why ToM impairments are rare [37]. Along with recent similar findings on the face network and mirroring network [37,59], we argue that these two features of the MTN white matter configuration—disproportionate local connectivity and redundant connections— commonly exist in social brain networks. Whether there are abnormalities in the amount or configuration of U-shaped fibers in individuals with autism spectrum disorder is not known but should be seriously examined in future research [76,77].

Third, we obtained a detailed and comprehensive picture of functional connectivity in the MTN. By examining coactivations across tasks that differentially tap ToM, we revealed an ‘intrinsic’ functional architecture that is constantly active and synchronized across contexts (Fig 3A). The coherence of this functional architecture, however, is a function of task demand. Prior studies showed that spontaneous ToM and effortful ToM recruit a common MTN but the latter elicits more regional activity [78]. Here we observed a similar effect on inter-regional connectivity––the greater degree of mentalizing a task requires, the stronger synchronization unfolds among MTN areas. As all network nodes were functionally defined by ToM task beforehand, this progressive effect on network-level coherence is most likely driven by mentalizing demand but less likely by other confounding factors (e.g. task difficulty or cognitive resource should affect global synchronization in dorsal attention network or frontoparietal control network, but not MTN). Indeed, additional analyses confirmed its specificity to mentalizing because no similar progressive effect can be observed in other brain networks (see Supplementary Fig 6; also see similar observations in [45]). In addition, we speculate that the hierarchical organization of functional connectivity may reflect the inherent order of information integration in this network (Supplementary Fig 3B), with posterior MTN areas processing low-level transient goals and differing perspectives while anterior MTN areas infer the meaning of social scenarios, and more temporally extended and contextually-deep traits and motivations. This serial-hierarchical information processing accords well with our current knowledge on the function of each MTN area [11] and converges with findings from recent MEG studies [79–81].

Our data allowed us to examine structure–function relationships in the MTN. Although structure leaves an indelible mark on function, a growing literature suggests that the link between SC and FC is complex, precluding a simple one-to-one mapping [82]. In other words, structural and functional networks do not necessarily have to be highly interdependent. Our results fully comply with this literature, showing that SC and FC were globally correlated across the entire network; however, they exhibited different network organization and fine-grained patterns. For example, some mentalizing areas ‘fired together’ but were not ‘wired together’ (e.g. TPJ-PreC and TPJ-DMPFC). In addition, the structural architecture was characterized by two parallel subsystems whereas the functional architecture featured a single hierarchical structure (Supplementary Fig 3B). Notably our results showed that the TPJ is a functional connectivity hub coordinating information integration within MTN, supporting a large body of prior research suggesting its pivotal role in ToM [4]. However, our results also uncovered an equally (or possible more) important role for the PreC. This region was not only a functional hub, but also the solo structural hub for the MTN. It is possible that the PreC plays a pivotal role in self-reflection/imagination/mental simulation processes that form the bedrock of high-level mentalizing [14].

Structure-function discrepancies were also prominent in other connectome features such as individual variance, spatial specificity, and functional specificity. We found that SC patterns were very similar across individuals, spatial extents, and DMN-vicinity networks, whereas functional connectivity and effective connectivity patterns were highly heterogeneous across conditions. Thus, the white matter architecture supporting mentalizing is consistent and stereotypical, indicating that the results of our SC analysis should readily generalize to different populations. By contrast, the MTN encompasses multiple functionally heterogeneous regions [83] and local voxels exhibit distinct functional selectivity and connectivity profiles (i.e. PreC [84], TPJ [85,86], ATL [87,88], DMPFC [89,90], VMPFC [91,92]). We are not unique in reporting this; other investigators have reported that FC is characterized by more variability than SC at the entire connectome level [59,93,94].

These connectome characteristics may have important methodological implications. The field of network neuroscience often considers brain networks as fixed and static entities [95] and conventionally defines network nodes by using atlas masks or meta-analytic coordinates. Our results suggest that a liberal selection of ROI locations might be workable for investigating the structural connectome but clearly this is problematic when measuring the functional connectome. Using group-level or Neurosynth coordinates to define MTN ROIs only preserves coarse patterns of SC but loses fine-grained fingerprints of FC and EC which reflect important dynamic and idiosyncratic aspects of the network (Fig 4). We urge future researchers to employ functional localizer and subject-specific ROIs when examining MTN connectivity, due to the disadvantages illuminated in this study of using liberal ROIs to define specific functional networks. In the same vein, precisely identifying individual-specific connectome would also be valuable for personalized psychiatry, for example when choosing brain stimulation sites to boost MTN dynamics and improve ToM performance in disorders with high heterogeneity such as autism and schizophrenia [96].

The key contribution of our work is in unraveling the relationship between the MTN and the default mode network. Previous meta-analyses and reviews have emphasized the strong spatial overlaps between the two networks [23,25–28,97,98]. It has been proposed that ToM is mediated by the DMN and thus the MTN is better characterized as a functional component of the DMN [24,28]. Indeed, the idea of ‘social cognition as the default mode of cognition’ is increasingly popular [24,99,100] and the DMN is believed to implement adaptive mentalizations that help individuals navigate their social environment by attributing mental states to others and spontaneously rehearsing social narratives to prepare for upcoming interactions [23,24,101]. To overcome the limitations of meta-analytic methods that rely on group-level analysis, the present study scrutinized the MTN-DMN relationship on a fine, single-subject scale (i.e. personalized MTN vs personalized DMN) and interrogated connectivity similarity rather than activation overlaps. We found that these networks are dissociable at multiple levels. First, the MTN and DMN exhibited similar patterns of SC but distinct patterns of FC and EC (Fig 5). Second, they showed the opposite network laterality. It is well known that regional activation in MTN is mostly right-lateralized (e.g. TPJ and ATL) [4,8,102–104]. Indeed, we found a right-hemisphere dominance at all levels of the mentalizing connectome (see Supplementary Table 3a). In contrast, our findings, as well as prior findings [105–107], found that the DMN was found to somewhat left-lateralized (Supplementary Table 3b). Last, we found brain connectivity patterns are relatively more homogeneous in the DMN, suggesting that the two systems have different degrees of inter-subject variation (Supplementary Table 6). Taken together, these findings dispute the prevalent assumption of identical network architecture between the MTN and DMN. They do share similar nodes and a similar wiring system but differ substantially in functional dynamics, laterality, and individual variability. Simply equating the two systems or fully adopting one’s network properties to another is problematic. While focusing on the commonality between the MTN and DMN has proven insightful in our understanding of fundamental brain mechanisms (e.g. ‘social by default’) [24,99,100], we contend that studying their differences is equally important and deserves further attention.

Our results shed light on the principles of DMN’s organization and function. The DMN has been proposed as a domain-general system for internal mental simulation [25,108]. It is recruited whenever people conjure up experiences outside of their local, immediate environment, such as thinking about the future or the past, mentally constructing places and spaces, and imagining hypothetical events and thinking about another’s perspective [109]. Here we found all DMN-vicinity networks were supported by a common anatomical architecture connecting major multisensory/multimodal areas (i.e. TPJ, ATJ, prefrontal cortices). This wiring configuration is essential for mental simulation because it enables simulation units to access to sensory inputs, stored conceptual knowledge, executive function resource, and evaluative processes. However, most DMN-vicinity networks exhibited distinct FC and EC patterns, indicating their functional roles for different simulation processing. These findings are compatible with the idea that the entirety of the DMN is responsible for mental simulation in general but different subnetworks are recruited for different forms and contents of simulation, such as constructing spatial and temporal models (episodic memory and prospection), narrative models (semantic memory), and social models related to self (self-referential), events (moral judgment) and other’s minds (ToM) [97,108,110]. These subnetworks are juxtaposed closely but communicate with different regions and networks outside the DMN for upstream and downstream processing of mental simulation [111]. Another possibility is that the DMN contributes to different types of mental simulation by forming distinct modes of connectivity that are distinguished by their location on the principle gradient of connectivity [112]. Future research should systematically study the functional specialization of subregions, subnetworks, gradients within the DMN and clarify their relationships with each domain-specific process [28,98,109,113–115].

Finally, the present work may prove useful for refining computational models of mentalizing. It is widely believed that ToM is a complex construct supported by multiple separate processes or computations [2]. Researchers have emphasized the role of different MTN areas in representing different representational and computational variables [7,18,43,116,117]. Much less is known about the relationship between these computations and how representational information is synthesized and transmitted within MTN [117]. Researchers could use our connectome features as priors into modelling of mentalizing operations, such as combining local representational information (i.e. RSA) with inter-regional connectivity properties (SC, FC, and EC), to simulate network dynamics more precisely and yield better behavioral predictions [118]. In addition, we hope our work catalyzes autism researchers to explore whether alterations in fine-grained connectivity explain deficits in mentalizing so prevalent in this disorder.

## Limitations

This study has some limitations. First, although we adopted a standard analytic pipeline, functional connectivity is inherently prone to conceptual and methodological problems [119] and is dependent on a variety of factors (e.g. ROI selection methods, task duration, motor demand, and stimuli features), thus findings should be interpreted cautiously [120–122]. Similarly, diffusion tractography has recently been criticized for having high false positives[123–125]. In addition, multimodal approaches could incur inflated structure-function discrepancies (e.g. PPI analyses might be a noisier or less reliable measure than the Pearson correlation and tractographies; or diffusion data is usually less smoothed than the functional data) [126]. Nevertheless, these tools provide considerable insights into the anatomical and functional architecture of the MTN. As both techniques develop, we hope other researchers replicate and extend our findings.

Second, as mentalizing is a broad concept, our conclusions were only based on the HCP ToM task, which might not be generalizable to other types of ToM tasks. We defined the spatial location of each MTN node by using peak activations in the social animation task and examined FC and EC in the same task. Although activation and connectivity were analytically orthogonal, this might introduce bias to some results (e.g. ToM-state FC was larger than emotion-state and motor-state FC in Fig 3A). In addition, there is massive heterogeneity in the tasks and neuroimaging methods used to investigate ToM [2,43]. The literature suggests that different ToM tasks (e.g. implicit or explicit) [9,127] and stimuli (e.g. false-belief stories, mind-in-the-eyes photos) might activate slightly different peak locations [9,11,17], which might result in different connectivity profiles. Future research should compare our results with connectivity patterns derived from other ToM tasks and other ROI selection methods to examine the impact of task format on connectome characteristics. Furthermore, since we were only interested in the core system of MTN (as well as the overlapped nodes between the MTN and DMN), some ToM-related regions (e.g. amygdala, inferior frontal gyrus, fusiform gyrus) and DMN regions (e.g. hippocampus, lateral temporal cortex, retrosplenial cortex) were not included in the present study. For simplicity, cross-hemispheric connections (e.g. between left and right TPJ) were also excluded in the analysis. Future research could add more MTN-related nodes and interhemispheric connections in the network analysis to further clarify the pathways and dynamics between core and extended MTN areas as well as across hemispheres.

Third, our investigation of the neural dynamics in MTN (e.g. PPI) is cursory and certain interpretations (e.g. hierarchical clustering results in Supplementary Fig 3B) are highly speculative. We used PPI because it is a simple static model of EC that provides some directional information about neural interactions (e.g. it builds up an explicit linear model of coupling between the seed and target region) [47–49,128,129]. However, we note that the post hoc interpretation of PPI results can be ambiguous as a significant increase in coupling from one region to another region may be significant when testing for a PPI in the opposite direction [48]. Dynamic causal modelling (DCM) can overcome these limitations but might not be easily incorporated in the present study. DCM with large sample size and model space (5 areas for 100 possible models) requires expensive computational resources [130] and the results (the optimal model) might not be generalizable to other ToM tasks (this is because different tasks have to set distinct sensory input areas (e.g. FFA for face-based ToM, VWFA for story-based ToM, and V5/MT for animation-based ToM [65]) and most of these input areas are beyond the core system of MTN). Future research should use more advanced EC methods [47,131–135], probably testing on a smaller fMRI dataset, to validate and extend our findings based on PPI.

Lastly, similar to other large-scale publicly available datasets, the HCP has inherent problems. In regards to this particular study, the social animation task was originally designed for children with autism [136] and is too easy for healthy adults, potentially result in ceiling effects for brain-behavior correlations (e.g. our connectome features can only explain less than 2% of variance). Future research needs to develop proper behavioral paradigms (e.g. using more ecologically valid social tasks) [137,138] to map individual ToM skills to connectome features and also test their reproducibility across multiple datasets [139].

## Conclusions

This multimodal neuroimaging study investigated the connectome basis of mentalizing processing. We found the anatomical architecture of the MTN is organized by two parallel subsystems (lateral vs medial) and constructed redundantly by local and long-range white matter fibers. We delineated an intrinsic functional architecture that is synchronized according to the degree of mentalizing and its hierarchy reflects the inherent information integration order in MTN. We examined the correspondence between the SC and FC and revealed their differences in network topology, individual variance, spatial specificity and functional specificity. Finally, we elaborated on the relationship between MTN and DMN. The two networks share similar nodes and SC but exhibited distinct FC patterns, the opposite laterality, and different homogeneity. In sum, we elucidated the structural and functional connectome supporting mentalizing processes and unraveled a complex relationship between the MTN and DMN. These findings have important implications on our understanding of how mentalizing is implemented in the brain and provide insights into the functions and organizations of the DMN.

## Materials and Methods

### Participants

All data used in the present study came from the WU-Minn HCP Consortium S900 Release. Subjects were included if they had completed all brain scans (T1/T2, tfMRI, rsfMRI, and dMRI). To reduce variance in the human connectome, we restricted our population to right-handed subjects, resulting in 680 healthy young adults in the final sample. It’s worth mentioning that only 672 out of 680 subjects were detected with enough robust signals in all five bilateral mentalizing ROIs in the functional ToM localizer task (see Supplementary Table 1), thus all findings in the present study were based on 672 subjects (379 females, 22-36 years old). No statistical methods were used to pre-determine this sample size but it is an order of magnitude larger than what is typically used in the field. The study was reviewed and approved by Temple University’s Institutional Review Board.

### Data Acquisition, Preprocessing and Analysis

Due to the complexity of the HCP data acquisition and preprocessing pipeline, listing all scanning protocols and data analysis procedures are beyond the scope of this paper; instead, they can be checked in full detail elsewhere [34,140–143]. Basically, we adopted the ‘minimally pre-processed’ volumetric images of task fMRI (tfMRI), resting-state fMRI (rsfMRI) and diffusion MRI (dMRI) that were provided by the HCP S900 release. The dMRI data had gone through EPI distortion, eddy current, and motion correction, gradient nonlinearity correction, and registration of the mean b0 volume to a native T1 volume. The fMRI data (rsfMRI and tfMRI) had undergone spatial artifact/distortion correction, cross-modal registration, and spatial normalization to MNI space. rsfMRI was further denoised using ICA-FIX. In addition to the HCP minimally pre-processed pipeline, we processed the dMRI data with FSL’s BEDPOSTX [35] to model white matter fiber orientations and crossing fibers, and denoised the tfMRI data with ICA-AROMA [144] to remove motion artifacts. All fMRI data were spatially smoothed at 4mm.

To ensure sensitivity to the connectome within each subject, we not only defined subject-specific ROIs based on the ToM localizer, but also performed all analyses firstly at the single subject level and then combined them into an aggregate statistic for group-level inference and significance tests. **Unless otherwise stated, all significant results reported in this study were corrected for multiple comparisons using false discovery rate (FDR).**

### Functional ToM Localizer and Selection of Mentalizing ROIs

The HCP social tfMRI data can be effectively used as a functional ToM localizer [142]. Participants were presented with short videos clips of geometric shapes either interacting in a socially meaningful way (e.g. dancing, coaxing, mocking, and seducing), or moving randomly. After watching each video, participants were required to choose between three possibilities: whether the moving shapes had a ‘Social interaction’, ‘No interaction’, or ‘Not sure’. Since the HCP S900 release had already provided the individual-level (within-subject) tfMRI analysis data (4mm smoothed MSM-All), we used the connectome workbench software to extract the MNI coordinates of the peak activation of five bilateral predefined MTN ROIs (as well as its magnitude). We used the contrast ‘ToM > Random’ for each subject at the individual level (see Supplementary Table 1). This contrast has been widely used to define mentalizing brain regions [2,11,142,145]. These subject-specific peak coordinates were used as input (6mm-radius spheres) in subsequent seed-based brain connectivity analyses at the individual level (probabilistic tractography, resting-state analysis, psychophysiological interaction), and the cluster peak magnitudes were adopted as the index of neural activity for mentalizing ROIs in brain-behavior association and hemispheric asymmetry analysis.

There are many ways to define the MTN. Different ToM tasks and paradigms activate slightly different sets of brain areas [10,11,17,20] but the core system (i.e. TPJ, PreC, ATL, DMPFC, VMPFC) is reliably and consistently implicated across tasks [11]. Since we were only interested in this core system, we did not include those extended regions that are only activated by certain task types, such as amygdala (i.e. affective processing), fusiform gyrus (i.e. agency/animacy detection) [146], inferior frontal gyrus (i.e. managing conflict between perspectives) [147]. In addition, based on a recent meta-analysis [11], we defined TPJ as a broad area encompassing both inferior parietal lobule (IPL, i.e. perspective taking) and posterior superior temporal sulcus (pSTS, i.e. biological motion perception). In that case, if TPJ had multiple clusters (which could fall either into traditional TPJ, or IPL or pSTS), we simply chose the strongest one to avoid excessive inter-subject inconsistency. For more detailed information about the mean and range of each ROI’s coordinates, please check Supplementary Table 1.

### ROI Definitions for DMN and Other Functional Networks

To define the nodes of multiple DMN-vicinity functional networks, we employed Neurosynth database to perform six automated meta-analyses of the functional neuroimaging literature on ‘mentalizing’, DMN, ‘autobiographical memory’, ‘self-referential’, ‘moral reasoning’ and ‘semantic memory’. For specific search terms queried in Neurosynth, please check Supplementary Table For each network, we extracted the cluster peak coordinates in the vicinity of TPJ, PreC, ATL, VMPFC, and DMPFC and used them as input (6mm-radius spheres) for subsequent seed-based brain connectivity analyses. For mental time travel, as there were no searchable terms in Neurosynth, we consulted the network node coordinates from a dedicated researcher-curated meta-analysis [148].

We also used a data-driven approach to define subject-specific DMN nodes. Spatial ICA was performed using the Group ICA of fMRI Toolbox (GIFT) [56] on HCP rsfMRI. A high model space (number of components =100) was estimated at the group level and we identified the ‘DMN-component’ (i.e. component 11, see Supplementary Fig 5) by visual inspection and spatial overlay with an independent DMN template (provided in GIFT) [149]. We decomposed the DMN template into ten binary masks (one for each DMN subregion) and used them separately to extract peak coordinates of the DMN-component at the single-subject level using subject-specific spatial ICA maps.

### Probabilistic Tractography

We used probabilistic tractography to reconstruct the entire MTN structural connectome. Tractography analyses were performed in native space and all results were transformed to Montreal Neurological Institute (MNI) standard space. We used an ROI-to-ROI approach where tractography was implemented between each pair of ROIs within the same hemisphere. Fiber tracking was initiated in both directions (from seed to target and vice versa) and 25000 streamlines were drawn from each voxel in the ROI. A binarized cerebellum mask was set as an exclusion mask for all analyses. The resulting 3D image files containing the output connectivity distribution were standardized using the maximum voxel intensity of each image resulting to a standardized 3D image with voxel values spanning from 0 to 1. These standardized path images were then thresholded at the 0.1 level to reduce false-positive fiber tracks. Binary connectivity maps were further generated for each subject and added across subjects.

FSL’s dtifit was used to fit a diffusion tensor model at each voxel. For each subject, the fractional anisotropy (FA), mean diffusivity (MD), radial diffusivity (AD), and axial diffusivity (RD) maps were created and their mean values for each ROI-ROI connection were extracted. The number of streamlines for each path was calculated by averaging two waytotal numbers produced by tractography. The connectivity probability for an ROI-ROI connection (e.g. TPJ-DMPFC) was defined as the streamline count of that connection divided by the sum of streamline counts of all connections passing either ROIs (e.g. there were 7 paths in total connecting either TPJ or DMPFC) [59].

### Community Detection, Hierarchical Clustering, and Graph Theoretical Analysis

To reveal the network organization of MTN, we performed three complementary analyses. We first prepared group-averaged connectivity maps for each hemisphere (i.e. using structural connectivity probability map or FC maps) and implemented graph theoretical analysis to reveal the hubness of the SC and FC using node centrality measures (see Supplementary Fig 3B for hubs of MTN, and Supplementary Table 5 for hubs of other DMN-vicinity networks). Next, we conducted community detection analysis on the same group-averaged connectivity maps to reveal the network modularity. Finally, we implemented agglomerative hierarchical clustering analysis to reveal the hierarchy of structural and functional connectome (Supplementary Fig 3B). For community detection analysis, modular partitions were obtained using Louvain community detection method in the NetworkX toolbox (https://networkx.github.io/). Graph theoretical analyses (node centrality) and agglomerative hierarchical clustering analyses were performed in MATLAB R2019b using function ‘centrality’ and ‘linkage’.

### Analyses of Major White Matter Bundles and Superficial White Matter System

For each hemisphere, ten major white matter bundles were identified for each subject using the Automated Fiber Quantification (AFQ) software package (https://github.com/jyeatman/AFQ) [38]. We focused our analysis on six major fiber tracts that were found to be critical for mentalizing processing in a recent meta-analysis [37]: the inferior longitudinal fasciculus (ILF), inferior fronto-occipital fasciculus (IFOF), cingulum (CING), uncinate fasciculus (UF), superior longitudinal fasciculus (SLF) and arcuate fasciculus (AF). We combined the results of SLF and AF because the AF is part of the SLF and their voxels overlap substantially in the AFQ [150]. We also analyzed four additional fasciculi provided by the AFQ (thalamic radiations, corticospinal tracts, anterior and posterior corpus callosum) and found little overlap (<1%) with all ROI-ROI connections, hence we did not include these tracts in the paper.

We used FSL’s atlasquery tool to evaluate the relative contribution of long-range and superficial white matter to the MTN [37,59]. After running probabilistic tractography, a binarized image for each ROI-ROI connection (standardized and thresholded at 0.1) was created for each subject and then added together across all subjects. We also combined all 20 bilateral connections together into one binarized image for each subject and added together across subjects to obtain the entire connectome image. Only voxels that existed in more than 50% of the subjects were retained (i.e. the skeleton image) and projected on two white matter atlases: the JHU white-matter tractography atlas with 48 long-range tract labels [41] and the LNAO superficial white matter atlas with 79 U-shaped bundles [42]. Voxelwise analyses were then implemented to calculate the probability of the skeleton image being a member of any labelled tracts within each atlas.

### Functional Connectivity and Effective Connectivity Analysis

To examine the coactivations and dynamics among MTN ROIs, we employed simple general linear regression model (GLM) to estimate the FC and EC during the ToM localizer task. For each MTN ROI as a seed region, we built a ‘generalized PPI’ model on HCP social tfMRI data with non-deconvolution method [151]. The model had 5 separate regressors: 2 psychological regressors of task events (ToM vs Random videos), 1 physiological regressor of the seed ROI’s timeseries, and 2 corresponding interaction regressors (task events x seed ROI’s timeseries). ToM-state FC between the seed ROI and other ROIs can be derived from the beta-weight of the physiological regressor (this was equivalent to the Pearson correlation between two ROIs’ timeseries after regressing out the task conditions and PPI regressors). The EC was estimated from the contrast between two interaction regressors (ToM PPI > Random PPI). Z-scored beta-weights were extracted for each pair of ROIs, which resulted in one 5×5 matrix for each connectivity type (ToM-state FC or EC), each hemisphere, and each subject. While the ToM-state FC matrices were further symmetrized, no symmetrization was applied to EC matrices due to its directional character. At the group level of the EC, one sample t-test across subjects was performed at each pair of ROIs to detect any significant effectivity connectivity. Non-parametric permutation tests (10,000 times) were also implemented to re-validate all significant results.

Coactivations among MTN ROIs were also examined in two other HCP tasks. The HCP motor task was originally designed to map motor-related areas. Participants had to follow visual cues to tap their left/right figures, squeeze left/right toes, or move their tongue [152]. Since it was inherently a non-social task, we used the motor task as a baseline check. The HCP emotion task was originally designed to elicit brain states for negative affect recognition and empathy, which are conceptually related to implicit mentalizing. Participants had to do a perceptual matching task either on emotional faces (fearful/angry) or neutral shapes [153].

Similar GLM models were employed to these two HCP tasks to derive task-state FCs among MTN ROIs. The model for motor task had 11 separate regressors: 5 psychological regressors of task events (left finger, right finger, left toe, right toe and tongue movement), 1 physiological regressor of the seed ROI’s timeseries, and 5 corresponding interaction regressors (task events x seed ROI’s timeseries). The model for HCP emotion task had 5 separate regressors: 2 psychological regressors of task events (emotional faces and neutral shapes), 1 physiological regressor of the seed ROI’s timeseries, and 2 corresponding interaction regressors (task events x seed ROI’s timeseries). Emotion-state FC between the seed ROI and other ROIs can be derived from the beta-weights of the physiological regressor.

Finally, we examined the coactivation patterns during resting state when mentalizing is assumed to be minimal. rsFC between five MTN ROIs was estimated by building five GLM models on HCP resting-state data. Each model defined one ROI’s time series as the dependent variable and the rest four MTN ROIs’ time series as independent variables. For each hemisphere, Fisher-transformed correlation coefficients (z-scored beta-weights) were extracted for each pair of 5×5 ROIs, symmetrized, and then averaged across two separate resting-state scans.

### Multilevel Modelling Analysis

Multilevel model analysis (MLM, also referred to as mixed effects regression models) was used to examine whether FC among MTN ROIs increased as the task category required more ToM processing. Task category was coded as 1, 2, 3 for HCP motor, emotion, ToM task, and was tested in MLM in SPSS 25.0 with random intercept to examine whether the degree of mentalizing required by a task category can predict the strength of FC for all ten pairwise connections.

### Structure-Function Relations

At the individual subject level, we prepared one connectivity matrix for SC, rsFC, ToM-state FC and EC, separately for each hemisphere. We did pairwise correlations among these brain connectivity matrices by taking all elements of each matrix except the diagonal ones (self-connections), applying a Fisher’s Z-transform, and then computing a Pearson correlation. Conventional one sample t-tests (against 0) were used at the group level to determine the statistical significance after controlling for multiple comparisons. Non-parametric permutation tests (10,000 times) were also implemented to re-validate all results.

### Brain-Behavior Associations

Pearson correlation was used to examine the brain-behavior association in SPSS 25.0. On the brain side, there were five types of features: (1) neural responses during mentalizing from 10 bilateral ROIs (the magnitude of BOLD signals in the contrast of ‘ToM > Random’); (2) white matter characteristics (i.e. connectivity probability, FD, MD, AD, RD, streamline counts) from 20 bilateral ROI-ROI connections and 20 major fasciculi; (3) 20 bilateral FC for resting state; (4) 20 bilateral FC for ToM-state; and (5) 40 bilateral EC for ToM-state. On the behavior side, we had two metrics from the HCP ToM task: task accuracy (Social_Task_TOM_Perc_TOM) and median reaction time (Social_Task_Median_RT_TOM). All significant results were corrected for multiple comparisons using false discovery rate (FDR).

### Hemisphere Lateralization

We used a 2-way repeated-measure ANOVA in SPSS 25.0 to examine MTN’s hemisphere lateralization at each level of measurements. At the neural activation level, we set the ANOVA with factors of ‘hemisphere’ and ‘5 MTN ROIs’. At the SC and rsFC level, we set the ANOVA with factors of ‘hemisphere’ and ‘10 ROI-ROI connections’. At the effective connectivity level, we set the ANOVA with factors of ‘hemisphere’ and ‘20 directional ROI-ROI connections’. If a main effect of hemisphere or an interaction effect was found in the ANOVA analysis, pairwise t-tests were further implemented to determine which hemisphere was dominant for which connections. Similar ANOVA was applied to DMN’s SC and rsFC to examine its hemispheric preference. All significant results were corrected for multiple comparisons using false discovery rate (FDR).

## Acknowledgments

We thank Dr. David V Smith for technical advice in PPI analyses and Dr. Axel Kohlmeyer for assistance with high performance cluster computing. We also thank 30 undergraduates from Temple University for their great work in inspecting all coordinates of mentalizing ROIs, especially Ying Lin, Richard Ho, Italia Hanik, Peyton Coleman, and Linda Jasmine Hoffman. The superficial white matter atlas (LNAO-SWM79) was generously provided by Dr. Pamela Guevara.

This work was supported by a Beijing Normal University start-up grant awarded to Y. Wang, a Temple University dissertation completion grant to A. Metoki, and National Institute of Health grants to I. Olson [RO1 MH091113 and R21 HD098509]. The content is solely the responsibility of the authors and does not necessarily represent the official views of the National Institutes of Health. The work also used Temple University’s High Performance Cluster Service (Owlsnest), which was supported by grants from National Science Foundation [#1625061] and the US Army Research Laboratory (W911NF-16-2-0189). The authors declare no competing financial interests.

HCP data were provided by the Human Connectome Project, WU-Minn Consortium (Principal Investigators: David Van Essen and Kamil Ugurbil; 1U54MH091657) funded by the 16 NIH Institutes and Centers that support the NIH Blueprint for Neuroscience Research; and by the McDonnell Center for Systems Neuroscience at Washington University.

## Supplementary Tables

**Supplementary Table 1.** Mentalizing ROIs. Mean and range of the subject-specific peaks for each mentalizing ROI identified in the HCP ToM task (with the contrast of ‘TOM > RANDOM’)

| MTN ROIs | MNI coordinates |  |  |  |  |  |  |  |  |
| --- | --- | --- | --- | --- | --- | --- | --- | --- | --- |
|  | Hemisphere | N | x | y | z | x-range | y-range | z-range | ES* |
| TPJ | R | 680 | -51 | -60 | 18 | 35-66 | -80 to -35 | 4-38 | 6.21 |
|  | L | 679 | 53 | -53 | 16 | -66 to -35 | -80 to -35 | 4-38 | 5.99 |
| ATL | R | 680 | 52 | -1 | -19 | 41-63 | -18 to 20 | -40 to -3 | 5.59 |
|  | L | 680 | -52 | -3 | -18 | -63 to -42 | -19 to 20 | -40 to -3 | 4.80 |
| PreC | R | 677 | 8 | -51 | 40 | 1-18 | -70 to -31 | 20-58 | 3.88 |
|  | L | 679 | -7 | -52 | 42 | -18 to 0 | -69 to -31 | 20-59 | 3.89 |
| DMPFC | R | 679 | 9 | 54 | 27 | 1-19 | 30-70 | 6-44 | 4.65 |
|  | L | 680 | -8 | 54 | 27 | -19 to 0 | 35-68 | 10-45 | 4.32 |
| VMPFC | R | 679 | 8 | 53 | -14 | 1-19 | 33-70 | -27 to 0 | 4.15 |
|  | L | 679 | -7 | 52 | -15 | -19 to -1 | 33-70 | -27 to 0 | 4.15 |

\*Effect size (ES) represents the mean signal change in the ROI (z-score from the peak voxel) across all subjects. At each individual level, any ROIs with weak responses to contrast 'TOM > RANDOM' (i.e. $ES < 1.5$ ) would be excluded for subsequent analyses. That's why some ROIs' N was smaller than 680. In sum, only 672 out of 680 subjects had all ten mentalizing ROIs.
TPJ: temporo-parietal junction; ATL: anterior temporal lobe; PreC: Precuneus; DMPFC: dorsomedial prefrontal cortex; VMPFC: ventromedial prefrontal cortex

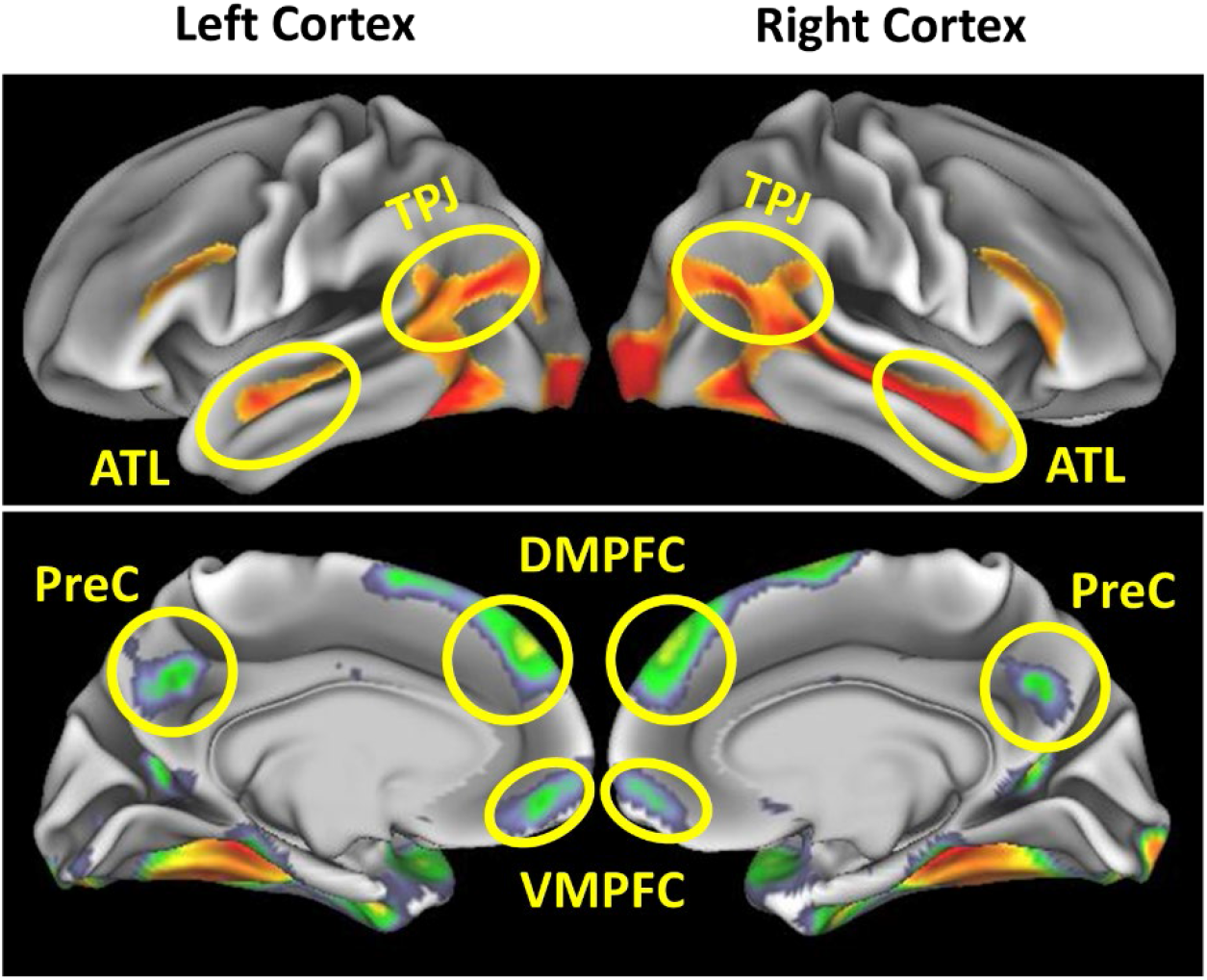
The spatial locations of each mentalizing ROI at the group-level. This group-level activation is only for illustration purpose. In fact, we independently defined each mentalizing ROI for each subject.

**Supplementary Table 2.** Functional connectivity (FC) across three HCP task-fMRI. **Table 2a.** Pairwise pattern similarity across three task categories

| Correlation of Task Pairs | Hemi-sphere | t | Sig. | Mean r | 95% CI |  | Effect size d |
| --- | --- | --- | --- | --- | --- | --- | --- |
| Motor FC & Emotion FC | LH | 26.49 | <.001 | 0.352 | 0.326 | 0.378 | 1.02 |
|  | RH | 26.85 | <.001 | 0.342 | 0.318 | 0.368 | 1.04 |
| Motor FC & ToM-state FC | LH | 37.71 | <.001 | 0.470 | 0.446 | 0.495 | 1.46 |
|  | RH | 37.60 | <.001 | 0.458 | 0.435 | 0.483 | 1.45 |
| Emotion FC & ToM-state FC | LH | 30.34 | <.001 | 0.391 | 0.366 | 0.416 | 1.17 |
|  | RH | 29.65 | <.001 | 0.375 | 0.350 | 0.400 | 1.14 |

**Table 2b.**
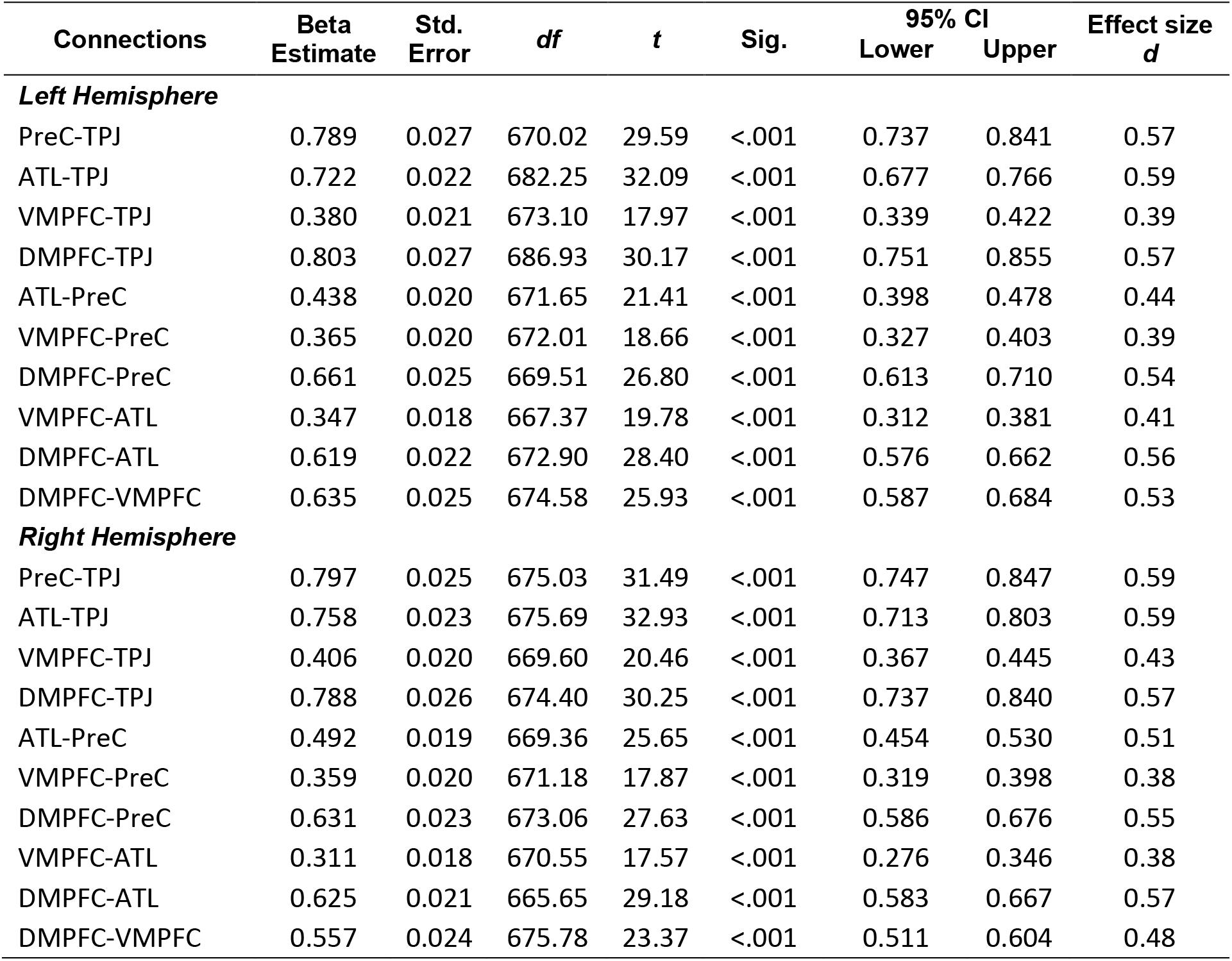
Results of multilevel models of FC across three task categories, indicating that FC in all 20 connections followed the same increasing trend from motor -> emotion -> ToM

**Supplementary Table 3.** Hemispheric asymmetry of the MTN and DMN at each level of measurements. **Table 3a:** The right-lateralized mentalizing network

|  | Right hemisphere<br>predominance | Left hemisphere<br>predominance |
| --- | --- | --- |
| **Neural**<br>**Activation** | ATL (t=15.32, p<0.001, d=0.59, 95% CI: 0.686, 0.887)<br>DMPFC (t=7.56, p<0.001, d=0.29, 95% CI: 0.249, 0.424)<br>TPJ (t=4.53, p<0.001, d=0.17, 95% CI: 0.129, 0.327) |  |
| **Structural**<br>**connectivity** | ATL-VMPFC (t=6.98, p<0.001, d=0.27, 95% CI: 0.153, 0.273) |  |
| **Resting-state**<br>**functional**<br>**connectivity** | TPJ-ATL (t=2.65, p=0.008, d=0.10, 95% CI: 0.010, 0.070)<br>PreC-VMPFC (t=2.86, p=0.004, d=0.11, 95% CI: 0.012, 0.066) | ATL-DMPFC (t=3.47, p=0.001, d=0.13, 95% CI: 0.020, 0.073) |
| **ToM-state**<br>**functional**<br>**connectivity** | TPJ-ATL (t=3.94, p<0.001, d=0.15, 95% CI: 0.138, 0.412) |  |
| **Effective**<br>**connectivity** | ATL→TPJ (t=5.05, p<0.001, d=0.20, 95% CI: 0.125, 0.284)<br>TPJ→ATL (t=4.89, p<0.001, d=0.19, 95% CI: 0.110, 0.258)<br>DMPFC→TPJ (t=3.27, p=0.001, d=0.13, 95% CI: 0.053, 0.211)<br>TPJ→DMPFC (t=3.11, p=0.002, d=0.12, 95% CI: 0.044, 0.196)<br>DMPFC→ATL (t=5.59, p<0.001, d=0.22, 95% CI: 0.131, 0.272)<br>ATL→DMPFC (t=5.93, p<0.001, d=0.23, 95% CI: 0.143, 0.284)<br>PreC→DMPFC (t=3.25, p=0.001, d=0.13, 95% CI: 0.046, 0.187)<br>VMPFC→DMPFC (t=2.92, p=0.004, d=0.11, 95% CI: 0.034, 0.176) |  |

**Table 3b:** The left-lateralized default mode network

|  | Right hemisphere<br>predominance | Left hemisphere<br>predominance |
| --- | --- | --- |
| **Structural**<br>**connectivity** | PreC-DMPFC (t=6.99, p<0.001, d= 0.27, 95% CI: 0.040, 0.071) | ATL-DMPFC (t=22.95, p<0.001, d= 0.89, 95% CI: 0.411, 0.488)<br>PreC-ATL (t=16.19, p<0.001, d= 0.62, 95% CI: 0.213, 0.272)<br>TPJ-ATL (t=25.92, p<0.001, d= 1.00, 95% CI: 0.397, 0.462)<br>ATL-VMPFC (t=8.26, p<0.001, d= 0.32, 95% CI: 0.146, 0.237)<br>TPJ-DMPFC (t=13.05, p<0.001, d= 0.50, 95% CI: 0.232, 0.315)<br>DMPFC-VMPFC (t=9.66, p<0.001, d= 0.37, 95% CI: 0.050, 0.075)<br>TPJ-PreC (t=4.17, p<0.001, d= 0.16, 95% CI: 0.046, 0.129)<br>PreC-VMPFC (t=11.94, p<0.001, d= 0.46, 95% CI: 0.066, 0.092)<br>TPJ-VMPFC (t=10.67, p<0.001, d= 0.41, 95% CI: 0.177, 0.257) |
| **Resting-**<br>**state**<br>**functional**<br>**connectivity** | TPJ-DMPFC (t=8.49, p<0.001, d= 0.33, 95% CI: 0.093, 0.149)<br>PreC-VMPFC (t=4.69, p<0.001, d= 0.18, 95% CI: 0.024, 0.057) | ATL-DMPFC (t=22.00, p<0.001, d= 0.85, 95% CI: 0.233, 0.278)<br>PreC-ATL (t=20.10, p<0.001, d= 0.78, 95% CI: 0.197, 0.240)<br>TPJ-ATL (t=22.12, p<0.001, d= 0.85, 95% CI: 0.301, 0.361)<br>ATL-VMPFC (t=17.60, p<0.001, d= 0.68, 95% CI: 0.154, 0.193)<br>PreC-DMPFC (t=7.85, p<0.001, d= 0.30, 95% CI: 0.060, 0.100)<br>TPJ-VMPFC (t=7.07, p<0.001, d= 0.27, 95% CI: 0.057, 0.101) |

**Supplementary Table 4.**
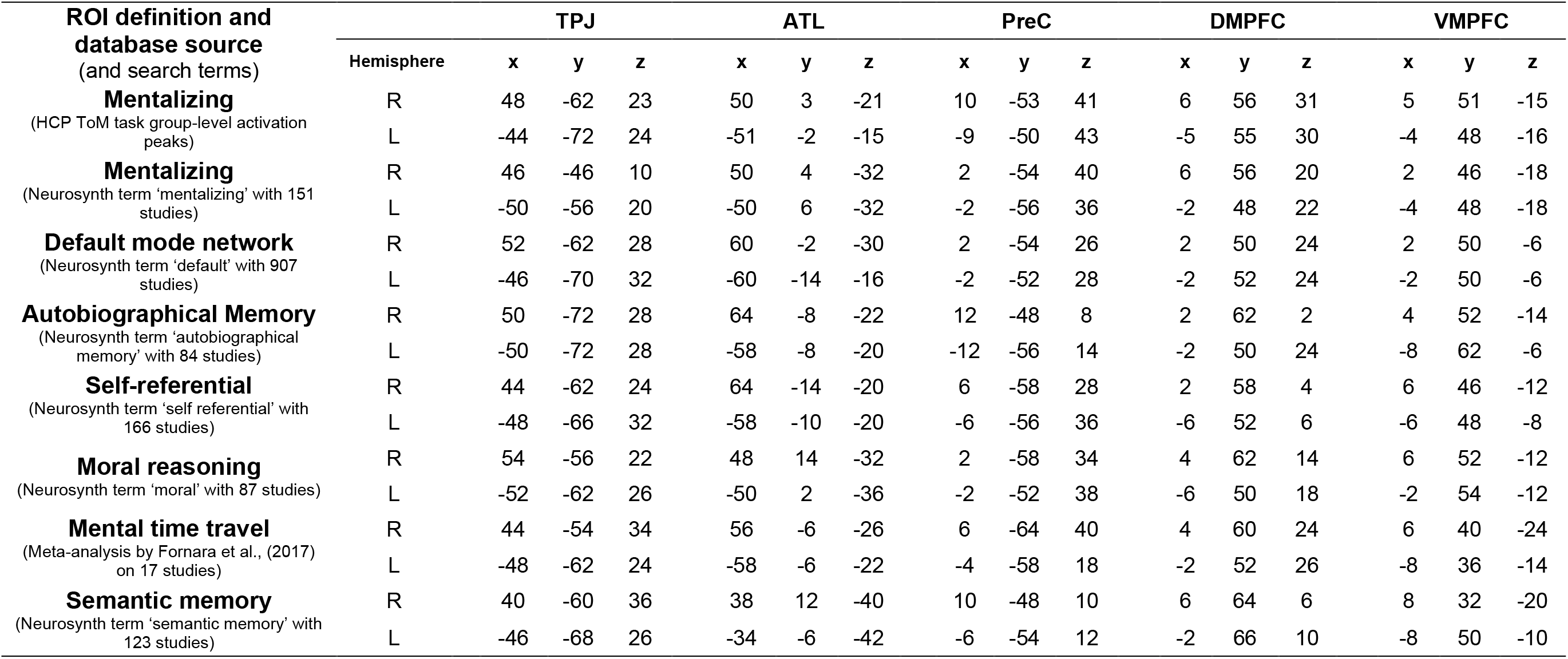
MNI coordinates of all functional ROIs used in the present study

**Supplementary Table 5.** Graph theoretic measure (node centrality) for each mentalizing ROI, showing its relative importance in the network. Asterisks and bold fonts indicate the hub.

|  | MTN | DMN | AMN | Self | Moral | MTT | SMN |
| --- | --- | --- | --- | --- | --- | --- | --- |
| **Structural Connectivity** |  |  |  |  |  |  |  |
| R-TPJ | 0.262 | 0.245 | 0.137 | 0.198 | 0.145 | 0.237 | 0.133 |
| R-ATL | 0.064 | 0.013 | 0.008 | 0.004 | 0.038 | 0.171 | 0.208 |
| R-PreC | **0.377\*** | **0.422\*** | **0.495\*** | **0.488\*** | **0.445\*** | **0.315\*** | **0.308\*** |
| R-DMPFC | 0.024 | 0.004 | 0.005 | 0.002 | 0.009 | 0.064 | 0.048 |
| R-VMPFC | 0.274 | 0.317 | 0.355 | 0.308 | 0.364 | 0.213 | **0.303\*** |
| L-TPJ | 0.260 | 0.260 | 0.178 | 0.202 | 0.231 | 0.238 | 0.147 |
| L-ATL | 0.055 | 0.012 | 0.055 | 0.004 | 0.037 | 0.043 | 0.013 |
| L-PreC | **0.383\*** | **0.440\*** | **0.416\*** | **0.478\*** | **0.386\*** | **0.406\*** | **0.478\*** |
| L-DMPFC | 0.026 | 0.005 | 0.030 | 0.001 | 0.010 | 0.014 | 0.010 |
| L-VMPFC | 0.276 | 0.284 | 0.321 | 0.315 | 0.337 | 0.299 | 0.352 |
| **Resting-state Functional Connectivity** |  |  |  |  |  |  |  |
| R-TPJ | **0.258\*** | **0.272\*** | 0.215 | 0.218 | **0.256\*** | **0.247\*** | **0.273\*** |
| R-ATL | 0.156 | 0.126 | 0.165 | 0.183 | 0.160 | 0.236 | 0.205 |
| R-PreC | **0.257\*** | 0.253 | **0.247\*** | **0.227\*** | 0.245 | 0.229 | 0.193 |
| R-DMPFC | 0.155 | 0.170 | 0.154 | 0.167 | 0.156 | 0.212 | 0.206 |
| R-VMPFC | 0.174 | 0.180 | 0.219 | 0.205 | 0.184 | 0.076 | 0.123 |
| L-TPJ | **0.259\*** | **0.254\*** | **0.281\*** | **0.293\*** | **0.301\*** | **0.313\*** | **0.324\*** |
| L-ATL | 0.155 | 0.162 | 0.128 | 0.121 | 0.115 | 0.155 | 0.090 |
| L-PreC | **0.256\*** | 0.233 | 0.251 | 0.281 | 0.260 | 0.228 | 0.233 |
| L-DMPFC | 0.168 | 0.187 | 0.189 | 0.146 | 0.160 | 0.204 | 0.143 |
| L-VMPFC | 0.162 | 0.164 | 0.150 | 0.159 | 0.163 | 0.100 | 0.210 |
| **ToM-state Functional Connectivity** |  |  |  |  |  |  |  |
| R-TPJ | **0.258\*** | **0.226\*** | 0.212 | 0.194 | **0.227\*** | **0.245\*** | **0.252\*** |
| R-ATL | 0.152 | 0.152 | 0.161 | 0.183 | 0.176 | 0.222 | 0.221 |
| R-PreC | **0.257\*** | **0.222\*** | **0.237\*** | **0.226\*** | **0.224\*** | 0.216 | 0.179 |
| R-DMPFC | 0.177 | 0.191 | 0.166 | 0.176 | 0.175 | 0.207 | 0.236 |
| R-VMPFC | 0.156 | 0.209 | 0.224 | 0.221* | 0.198 | 0.109 | 0.112 |
| L-TPJ | **0.250\*** | **0.230\*** | **0.256\*** | **0.250\*** | **0.256\*** | **0.268\*** | **0.265\*** |
| L-ATL | 0.150 | 0.168 | 0.154 | 0.153 | 0.151 | 0.187 | 0.119 |
| L-PreC | **0.248\*** | 0.219 | 0.219 | **0.246\*** | 0.224 | 0.191 | 0.231 |
| L-DMPFC | 0.186 | 0.194 | 0.198 | 0.171 | 0.190 | 0.233 | 0.164 |
| L-VMPFC | 0.166 | 0.189 | 0.172 | 0.180 | 0.178 | 0.122 | 0.220 |

Abbreviation: MTN: mentalizing network; DMN: default-mode network; AMN: autobiographical memory network; MTT: mental time travel; SMN: semantic memory network;

**Supplementary Table 6.** Comparison of individual variability between the MTN and DMN.

|  |  | **SC** | **rsFC** | **ToM-FC** | **EC** |
| --- | --- | --- | --- | --- | --- |
| Mean group-to-individual similarity $r$ (sd, [95%CI]) | | | | | |
| **MTN** | L | 0.76 (0.21, [0.74,0.78]) | 0.43 (0.30, [0.40,0.45]) | 0.42 (0.31, [0.40,0.45]) | 0.14 (0.31, [0.11,0.16]) |
|  | R | 0.77 (0.21, [0.75,0.78]) | 0.43 (0.29, [0.41,0.45]) | 0.44 (0.33, [0.42,0.47]) | 0.14 (0.34, [0.11,0.16]) |
| **DMN** | L | 0.92 (0.08, [0.91,0.93]) | 0.53 (0.31, [0.50,0.55]) | 0.44 (0.34, [0.38,0.49]) | 0.15 (0.30, [0.11,0.18]) |
|  | R | 0.91 (0.08, [0.90,0.92]) | 0.70 (0.20, [0.69,0.72]) | 0.59 (0.30, [0.54,0.64]) | 0.15 (0.29, [0.11,0.18]) |

**Supplementary Table 7.** Summary of all methods, analyses and findings in the present study

|  | Goals | Data | Analyses | Methods | Findings |
| --- | --- | --- | --- | --- | --- |
| **Network nodes selection** | to define subject-specific MTN ROIs | HCP ToM tfMRI | localize peak activation from the contrast 'ToM > random' in each subject | peak coordinates extraction for five bilateral MTN ROIs (Connectome workbench) |  |
|  | to define putative DMN ROIs | Neurosynth | extract node coordinates from Neurosynth meta-analytic maps | Search term 'default' in Neurosynth database |  |
|  | to define subject-specific DMN ROIs | HCP rsfMRI | extract peak coordinates of the DMN component in each subject | Spatial ICA (GIFT toolbox) |  |
|  | to define five DMN-adjacency networks | Neurosynth | extract node coordinates from Neurosynth meta-analytic maps | Search terms can be found in Supplementary Table 4 |  |
| **Structural Connectome** | to reveal structural architecture of the MTN | HCP dMRI | reconstruct MTN connectome by running ROI-to-ROI tractography between MTN ROIs | probabilistic tracking, community detection, hierarchical clustering | two parallel subsystems (medial vs lateral) |
|  | to reveal important major fasciculi in a quantitative way | HCP dMRI | reconstruct ten major fasciculi and overlap with the MTN connectome | AFQ toolbox | five key fasciculi for the MTN |
|  | to reveal relative contributions of short-range and long-range fibers to the MTN | HCP dMRI | calculate the overlap between MTN connectome and long-range and superficial fiber systems | FSL Atlasquery | MTN was redundantly supported by long and superficial fibers, but primarily by the latter |
| **Functional Connectome** | to reveal intrinsic functional architecture of the MTN | HCP rsfMRI; Motor, Emotion, ToM tfMRI | calculate FC between MTN ROIs across four HCP tasks (resting, motor, emotion, and ToM) | GLM models, multilevel modeling, community detection, hierarchical clustering | FC had a hierarchical structure, and was increased when the task requires more mentalizing. |
|  | to reveal task-modulated dynamics during mentalizing | HCP ToM tfMRI | build simple EC (PPI) models between MTN ROIs | FSL gPPI | EC exhibited strong right hemisphere predominance during mentalizing |
| **Individual variance** | to reveal individual variability of the MTN connectome | HCP dMRI, rsfMRI, ToM tfMRI | calculate group-to-individual and subgroup-to-subgroup similarity for SC, FC, and EC | split-half cross-validation<br>leave-one-out cross-validation | large homogeneity in SC, medium in FC, and small in EC. |
| **Spatial specificity** | to explore how MTN connectivity patterns are spread & changed in space | HCP dMRI, rsfMRI, ToM tfMRI | compare connectivity patterns between subject-specific MTN, group-level MTN, Neurosynth MTN, and shuffled-ROI MTN | pattern similarity analysis | FC and EC patterns were highly heterogeneous across space whereas SC patterns remained similar. |
| **Functional specificity** | to examine the connectome resemblance between the MTN and DMN | HCP dMRI, rsfMRI, ToM tfMRI | compare SC, FC, EC patterns between MTN, DMN, and other five DMN-adjacency networks | pattern similarity analysis<br>graph theoretical analysis | MTN and DMN had similar SC, but distinct FC and EC patterns, and opposite lateralization. |
| **Structure-function relation** | to examine SC-FC correspondence within the MTN | HCP dMRI, rsfMRI, ToM tfMRI | compare connectivity patterns and network organization (hubness, topology) between SC, FC and EC | pattern similarity analysis<br>graph theoretical analysis | SC and FC were tightly linked but had different fine-grain patterns and underlying architecture (parallel vs hierarchical) |
| **Lateralization** | to reveal hemispheric asymmetry in the MTN and DMN | HCP dMRI, rsfMRI, ToM tfMRI | compare task-evoked BOLD signals, SC, FC, EC across two hemispheres | SPSS ANOVA and t-tests | The MTN was right-lateralized whereas the DMN was left-lateralized |
| **Brain-behavior association** | to explore if any connectome features were associated with ToM task performance | HCP dMRI, rsfMRI, ToM tfMRI | run correlations between MTN features (task responses, SC, FC, EC) and ToM task performance (accuracy and speed) | SPSS Pearson correlations | Task responses in right ATL and TPJ as well as their EC were correlated with ToM task accuracy |

## Supplementary Figures

**Supplementary Figure 1.**
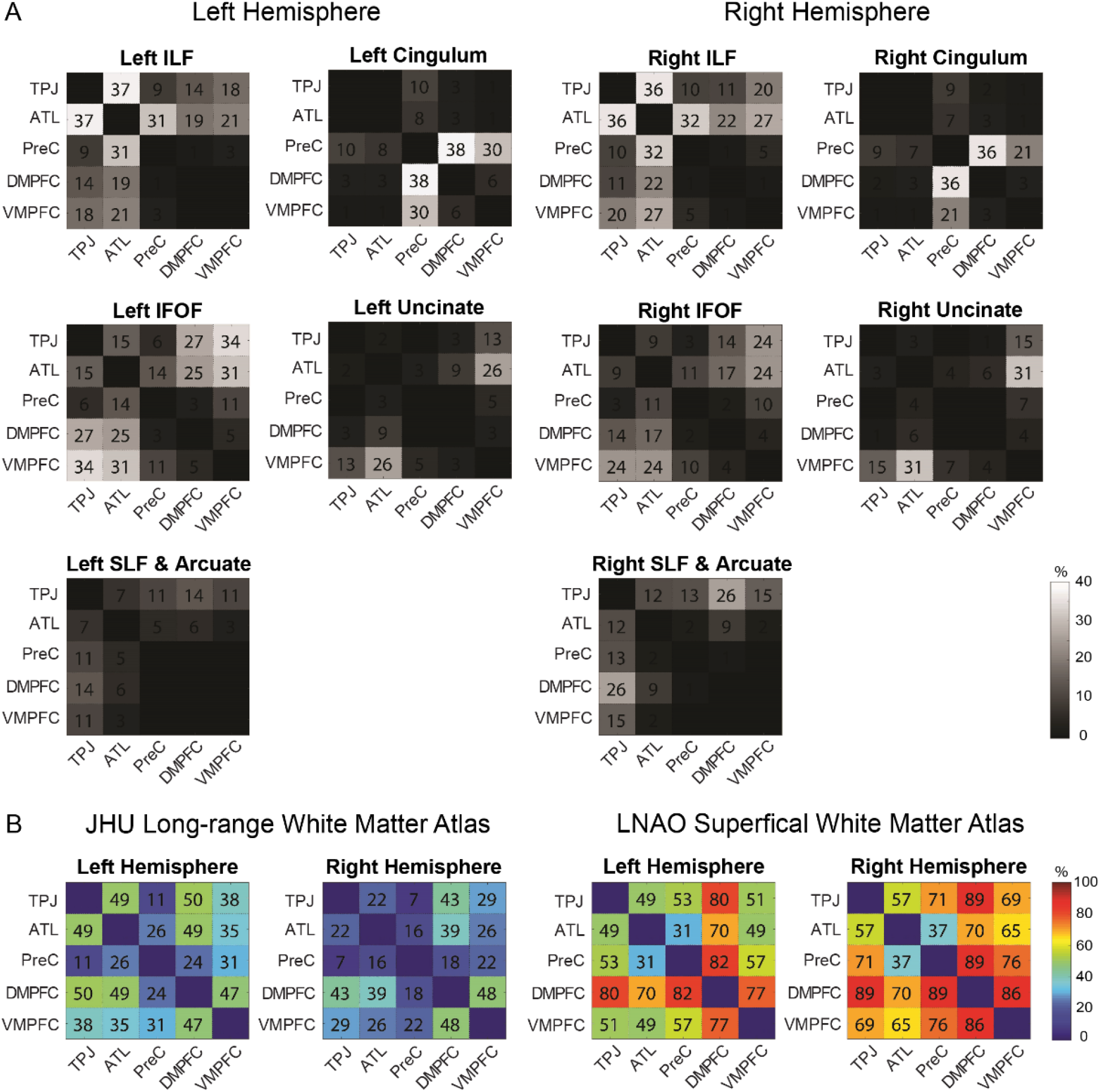
(A) The overlap between major white matter fasciculi and the mentalizing connectome in each hemisphere. Each number in the matrices represents the mean overlapped trajectories between an ROI-ROI tract and a major fasciculus across all subjects. (B) The relative contributions of long-range vs superficial white matter to the mentalizing connectome in each hemisphere. For most ROI-ROI connections, they were mediated more by the superficial white matter system than the long-range fiber system. Abbreviations: inferior longitudinal fasciculus (ILF), inferior fronto-occipital fasciculus (IFOF), superior longitudinal fasciculus (SLF)

**Supplementary Figure 2.**
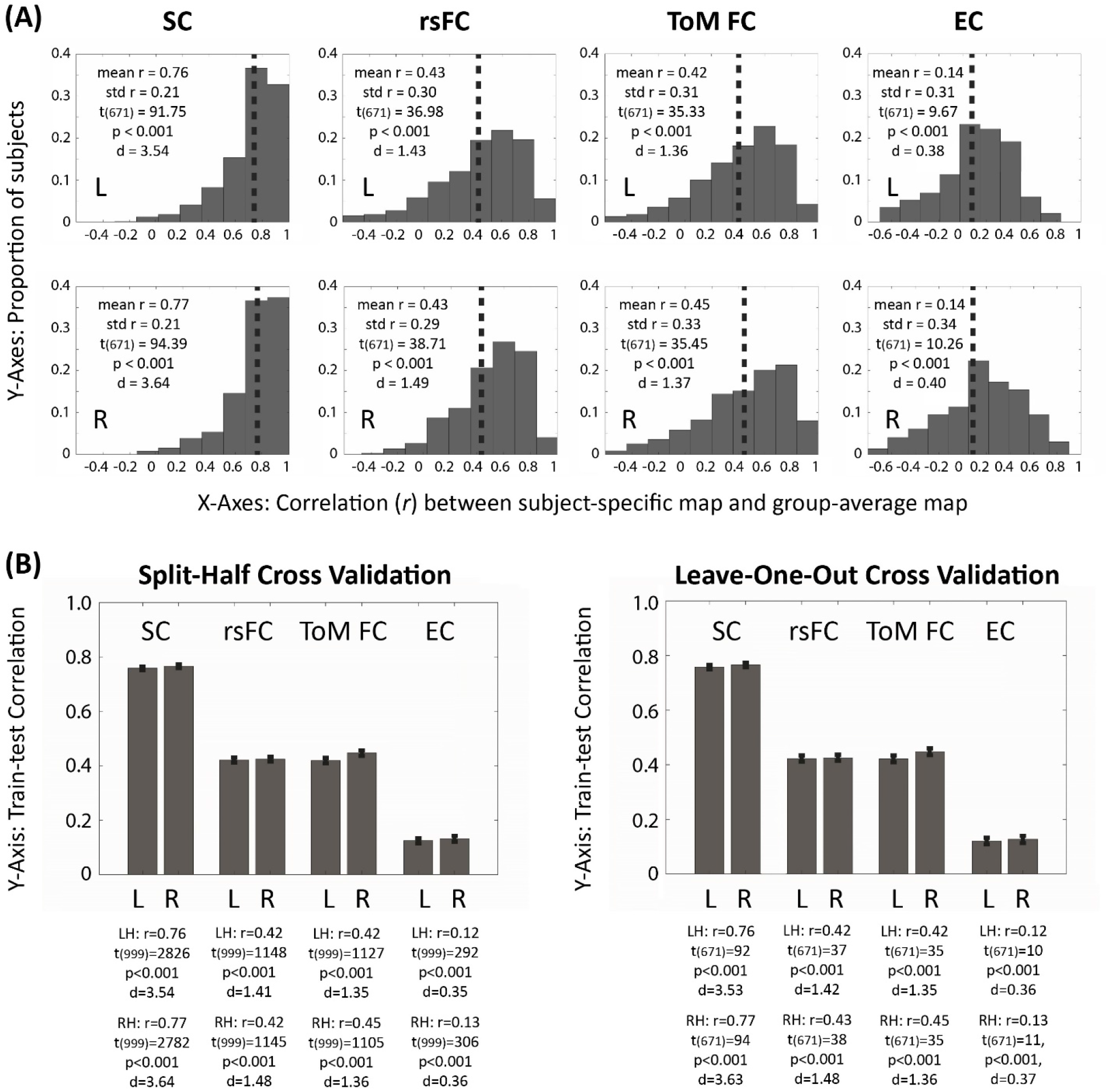
Individual Variance of the Mentalizing Connectome. (A) For each subject, we calculated the similarity between group-averaged brain connectivity matrices and subject-specific ones. Histograms indicated that most subjects exhibited high group-to-individual similarity for the SC, rsFC and ToM-state FC (52.3% subjects had *r*>0.5) and mild group-to-individual similarity for the EC (57.2% subjects had *r* >0.1). Dash line indicates mean group-to-individual correlation. (B) Two cross-validation (CV) analyses were further performed to measure group-to-group and group-to-individual similarity. The split-half CV randomly selected 50% subjects and used their averaged matrices to predict each of the rest 50% subject’s matrices. This train-test procedure was repeated for 1000 times to calculate the overall subgroup-to-subgroup similarity. The leave-one-out CV (LOOCV) built group-averaged matrices from n-1 subjects and used them to predict the matrices in a new subject. This LOOCV train-test procedure was repeated for all 672 subjects to calculate the overall group-to-individual similarity. Consistent with the correlation distribution in (A), both CV schemes found very large train-test correlation for the SC, medium correlation for rsFC and ToM-FC, and small correlation for the EC. Error bar indicates standard error (SE).

**Supplementary Figure 3.**
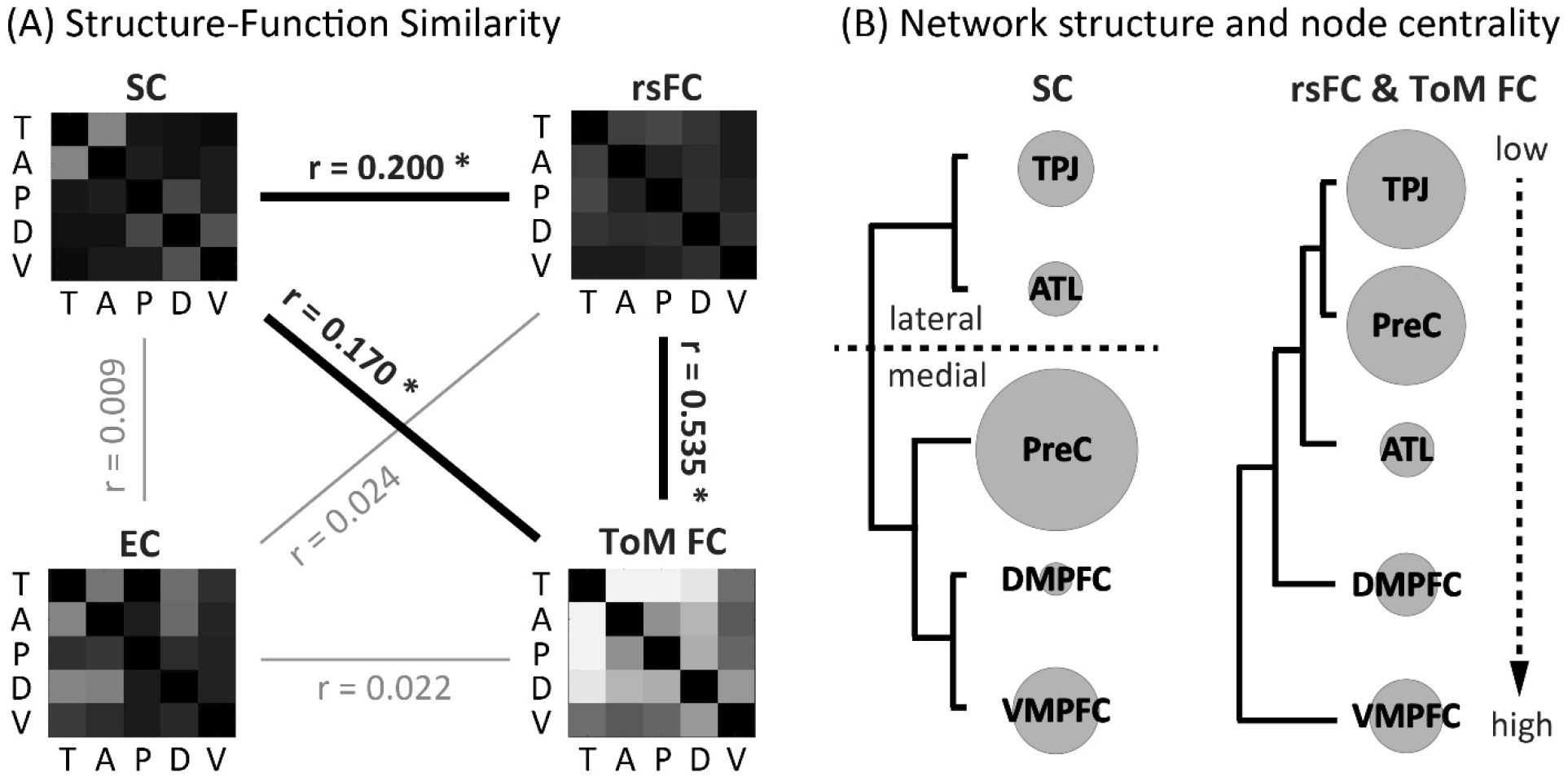
Structure-Function Relation and Network Architecture. (A) We explored the structure-function relationship in the MTN by examining the correspondence between FC, rsFC, ToM-FC and EC. Results suggest that the SC, rsFC, and ToM-state FC were tightly linked but EC was not correlated with any of them. Although the global patterns of SC and FC were largely matched, SC-FC disagreement can still be observed in certain connections (i.e. PreC-TPJ and DMPFC-TPJ had strong FC but weak rsFC and ToM-state FC). (B) We used hierarchical clustering and graph theory analyses to reveal the organization of structural and functional connectome. Results suggest that the structural connectome was constructed by two parallel subsystems (medial vs lateral) and precuneus was the structural hub, whereas the functional connectome was organized as a serial-hierarchical structure with TPJ and Precuneus as two functional hubs. As a post-hoc interpretation, we speculate that this hierarchical organization indirectly reflect the information integration order during mentalizing, as it accords well with the literature summarizing the specific function of each MTN area. Taking the social animation task as an example, the TPJ and PreC work together to infer low-level transient mental states by mentally simulating the context and taking perspectives of each geometric character (e.g. ‘the big triangle wants to catch the small triangle’); the ATL subsequently retrieves social semantic knowledge so that one can conceptually understand the scenario (e.g. ‘this is a chasing behavior’). Lastly, the DMPFC is engaged in making high-level inferences such as ascribing enduring traits to each character (e.g. ‘the big triangle is powerful, but the small triangle is agile’), and portions of the VMPFC may contribute to the overall emotional and motivational evaluation (e.g. ‘they happily play together’). The size of the gray circle is scaled to reflect the node centrality (see Supplementary Table 5). For (A) and (B), only results in right hemisphere is showed here but the left hemisphere has very similar findings. Abbreviations: T=Temporo-parietal junction (TPJ); A=Anterior Temporal Lobe (ATL); P=Precuneus (PreC); D=Dorsal Medial Prefrontal Cortex (DMPFC); V=Ventral Medial Prefrontal Cortex (VMPFC)

**Supplementary Figure 4.**
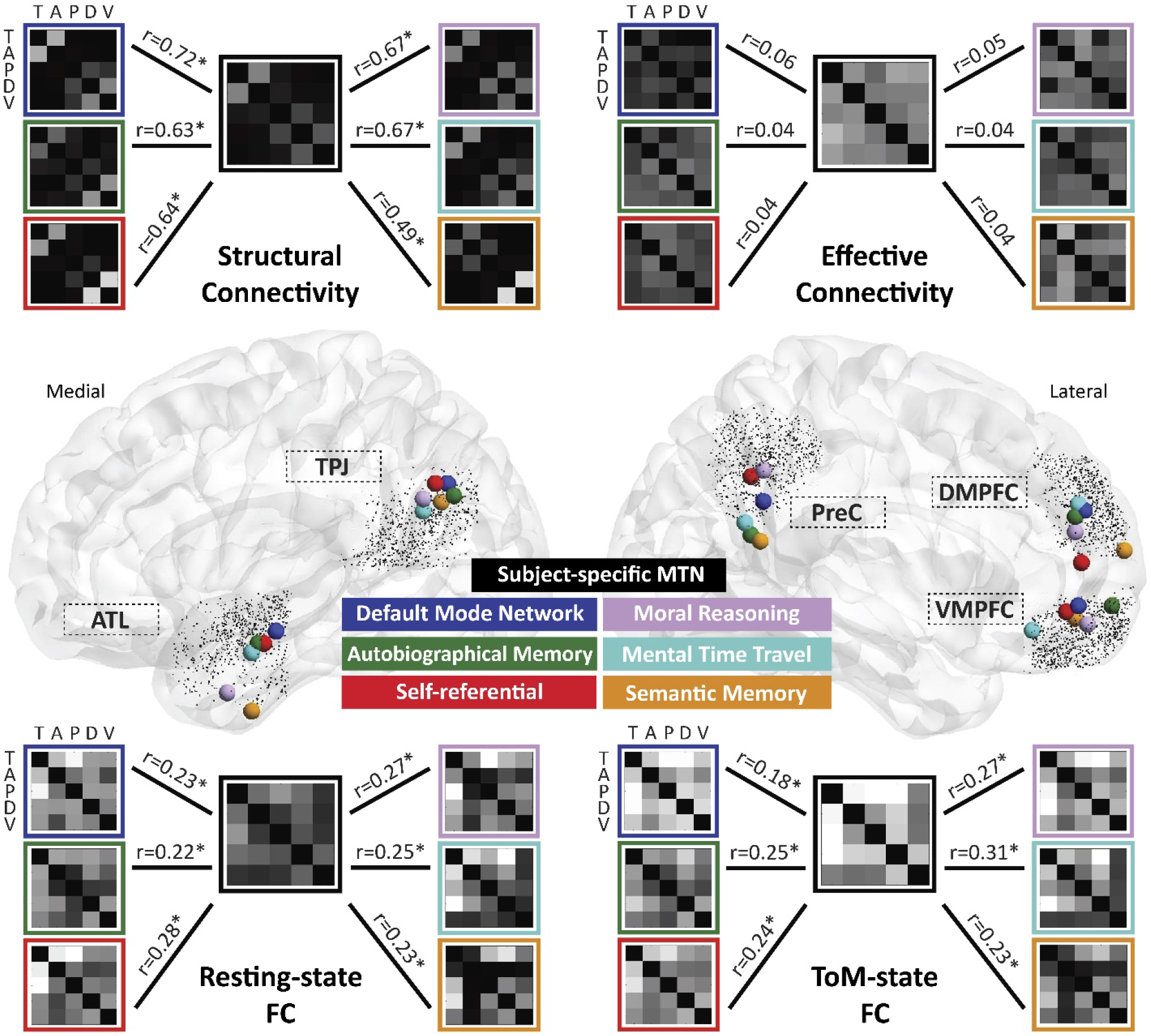
Brain connectivity similarity between MTN, DMN and other DMN-vicinity networks in the left hemisphere. Asterisks indicate statistical significance (p<0.05). Abbreviations: T=Temporo-parietal junction (TPJ); A=Anterior Temporal Lobe (ATL); P=Precuneus (PreC); D=Dorsal Medial Prefrontal Cortex (DMPFC); V=Ventral Medial Prefrontal Cortex (VMPFC)

**Supplementary Figure 5.**
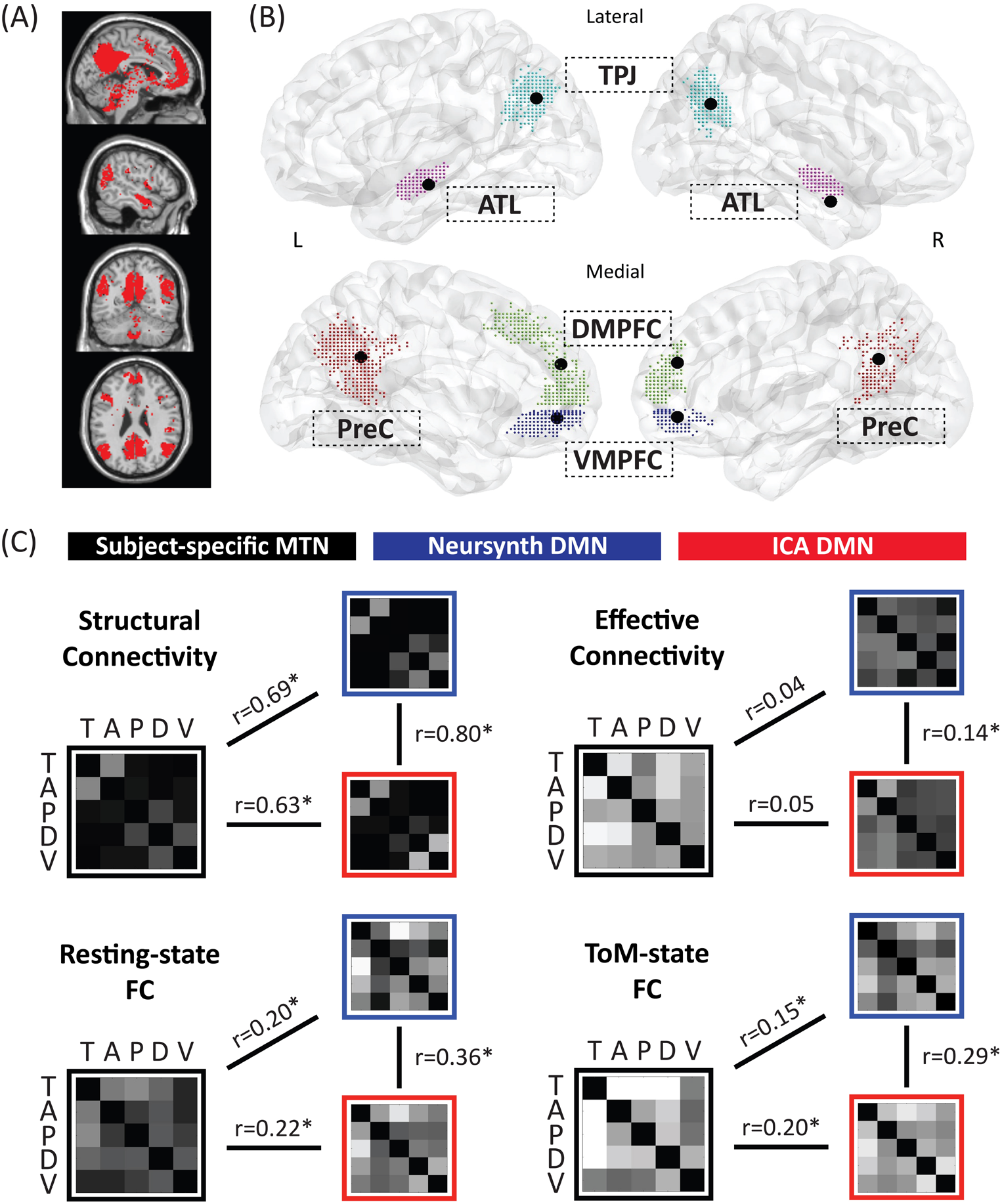
Brain connectivity similarity between subject-specific MTN, Neurosynth-DMN and subject-specific ICA-DMN. (A) We performed group ICA across all subjects and the ‘DMN-component’ was identified (i.e. component 11 was showed here in red). (B) We then extracted peak coordinates of the DMN-component at single-subject level using subject-specific spatial ICA maps (each dot indicates a subject’s location). (C) We found the subject-specific ICA-DMN and Neurosynth-DMN exhibited the same relationship with the subject-specific MTN, with highly similar structural connectivity but different patterns of FC and EC. Asterisks indicate statistical significance of Pearson correlation between two patterns (p<0.05). Abbreviations: T=Temporo-parietal junction (TPJ); A=Anterior Temporal Lobe (ATL); P=Precuneus (PreC); D=Dorsal Medial Prefrontal Cortex (DMPFC); V=Ventral Medial Prefrontal Cortex (VMPFC)

**Supplementary Figure 6.**
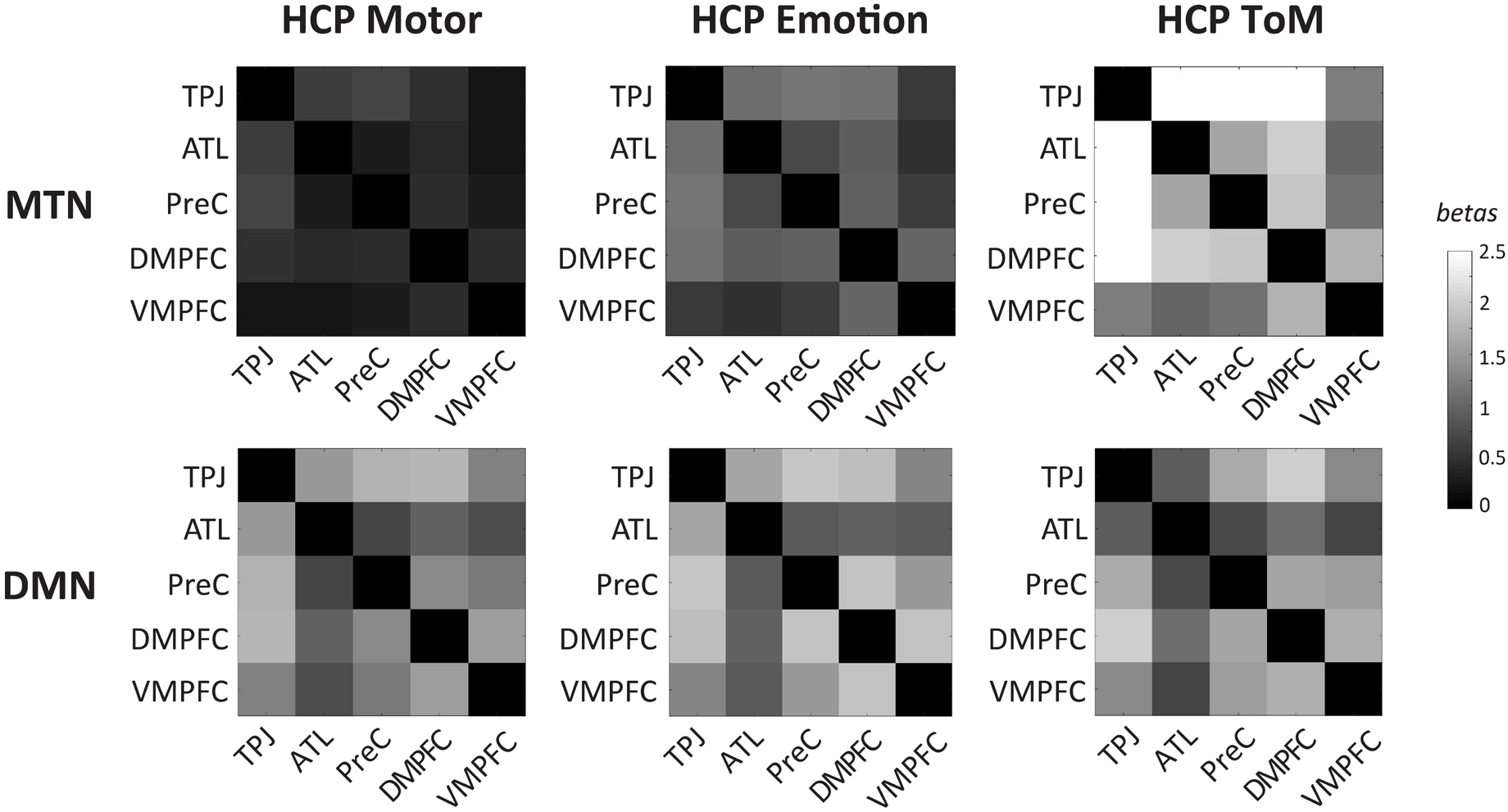
Functional connectivity (FC) in the MTN and DMN across different cognitive tasks. In the main text Fig 3A, we showed that the MTN had a progressively increased network-level FC from the motor-state, emotion-state, to the ToM-state task (as showed here in the first row). To further examine whether this effect was specific to the MTN or any brain networks, we performed analogous cross-tasks analyses on the DMN (see the second row). Similar to the MTN, we found a consistent FC topological patterns across three HCP task in the DMN, which indicates the intrinsic coherent functional architecture of the DMN; however, we did not observe the progressive FC trend existed in the DMN, suggesting that the DMN FC remained similar across three cognitive states. This finding is consistent with a previous study using the same HCP dataset (see Figure 6 in Cole et al., 2014, Neuron [45]). For completeness, we also examined another no-mentalizing task (gambling task) and moderate-mentalizing-demand task (face localizer) in the HCP dataset and got consistent and predicable findings (i.e. no-mentalizing = weak FC; moderate mentalizing = medium FC). For simplicity and readability, we did not show their results here. In sum. this additional analysis confirms that the progressive effect is exclusive to the MTN and supports our claims that the greater degree of mentalizing a task requires, the stronger synchronization unfolds among MTN areas.

